# Mechanisms of ventricular arrhythmias elicited by coexistence of multiple electrophysiological remodeling in ischemia: a simulation study

**DOI:** 10.1101/2021.08.24.457584

**Authors:** Cuiping Liang, Qince Li, Kuanquan Wang, Yimei Du, Wei Wang, Henggui Zhang

## Abstract

Myocardial ischemia, injury and infarction (MI) are the three stages of acute coronary syndrome (ACS). In the past two decades, a great number of studies focused on myocardial ischemia and MI individually, and showed that the occurrence of reentrant arrhythmias is often associated with myocardial ischemia or MI. However, arrhythmogenic mechanisms in the tissue with various degrees of remodeling in the ischemic heart have not been fully understood. In this study, biophysical detailed single-cell models of ischemia 1a, 1b, and MI were developed to mimic the electrophysiological remodeling at different stages of ACS. 2D tissue models with different distributions of ischemia and MI were constructed to investigate the mechanisms of initiation of reentrant waves during the progression of ischemia. Simulation results in 2D tissues showed that the vulnerable window (VW) in the tissue with multiple pathological conditions were determined by the VWs in the tissues with a single pathological condition. In different pathological conditions, action potential duration (APD) and the conduction velocity (CV) change differently. In the tissue with multiple pathological conditions, when the borders of different pathological conditions were perpendicular to the excitation wavefront, reentrant waves were mainly induced by the spatial heterogeneity of refractory periods due to the interaction of APD and CV along the wavefront. When the borders were parallel to the wavefront, the increased excitation threshold of MI region as well as the impaired excitability of ischemia region were the primary reason for the generation of reentry. Finally, the reentrant wave was observed in a 3D model with a scar reconstructed from MRI images of a MI patient. These comprehensive findings provide novel insights for understanding the arrhythmic risk during the progression of myocardial ischemia and highlight the importance of the multiple pathological stage in designing medical therapies for arrhythmia in ischemia.

**Author summary:** Abnormal initiation or conduction of electrical impulses may lead to cardiac arrhythmias, which are very important cause of sudden and early death in developed countries. In many cases, cardiac arrhythmias are accompanied by ventricular fibrillation sustained by re-entry. The occurrence of reentrant arrhythmias is often associated with acute coronary syndrome, including three phases of myocardial ischemia 1a, 1b, and infarction. Previous studies have made lots of efforts to unravel the mechanisms of the initiation and maintenance of reentry waves during myocardial ischemia or infarction. However, the mechanisms of initiation of reentrant waves in the tissue with multiple ischemic remodeling are not fully understood. Multi-scale computational models at the cellular, tissue, and organ levels were developed in this study. The main finding in this study is that reentrant waves in multiple pathological tissues were mainly induced by the spatial heterogeneity of refractory periods, when the borders of different pathological conditions were perpendicular to the wavefront of the excitation wave. In addition, the increased excitation threshold of MI region as well as the impaired excitability of ischemia region can induce the generation of reentry, when the borders were parallel to the excitation wavefront. This provides insights into mechanisms of ventricular arrhythmias elicited by coexistence of multiple electrophysiological remodeling in ischemia.

## Introduction

During the progression of coronary artery stenosis or ACS, differential electrophysiological remodeling in ion currents was observed. For example, in the stage 1a of ischemia (vascular obstruction <15min), I_Na_ decreased and activation delayed, I_NaL_ increased and I_CaL_ decreased, resulting in the impairment of the excitability of cell and the decrease of APD. In the stage 1b of ischemia (vascular obstruction 15∼45 min), I_CaL_ further decreased, and I_NaCa_ and I_NaK_ were also inhibited, resulting in the further decrease of APD. The long-lasting ischemia (vascular obstruction up to 3∼5days; referred as MI stage in this study) produced the largest APD in these three phases due to the decline of I_Kr_ and I_Ks_, although I_Na_ and I_CaL_ were still inhibited to some extent in MI (see Table 1, Fig 2, and S1 Fig for details). At a certain stage, differential electrophysiological remodeling can coexist in the same heart tissues[1, 2]. Previous studies have put numerous efforts to unravel the mechanisms responsible for the initiation and maintenance of reentry waves at both single-cell level and tissue level under the condition of single electrophysiological remodeling. For example, for stage of ischemia 1a, Kazbano *et al.* studied the effects of each single component (hyperkalemia, hypoxia, and acidosis) of ischemia on reentry wave propagation[3]. In addition, the inhomogeneous distribution of ion currents[4] and cross wall heterogeneity[5] in ischemia 1a will enhance the vulnerability to reentry. For the MI stage (i.e., the long-lasting ischemia), the changes of electrophysiological characteristics and the increased heterogeneous distribution of scar tissues will facilitate the generation and maintenance of reentrant waves[6, 7]. However, the combined effect of multiple myocardial ischemic stages on the development of reentrant arrhythmias is rarely investigated.

**Table 1.**
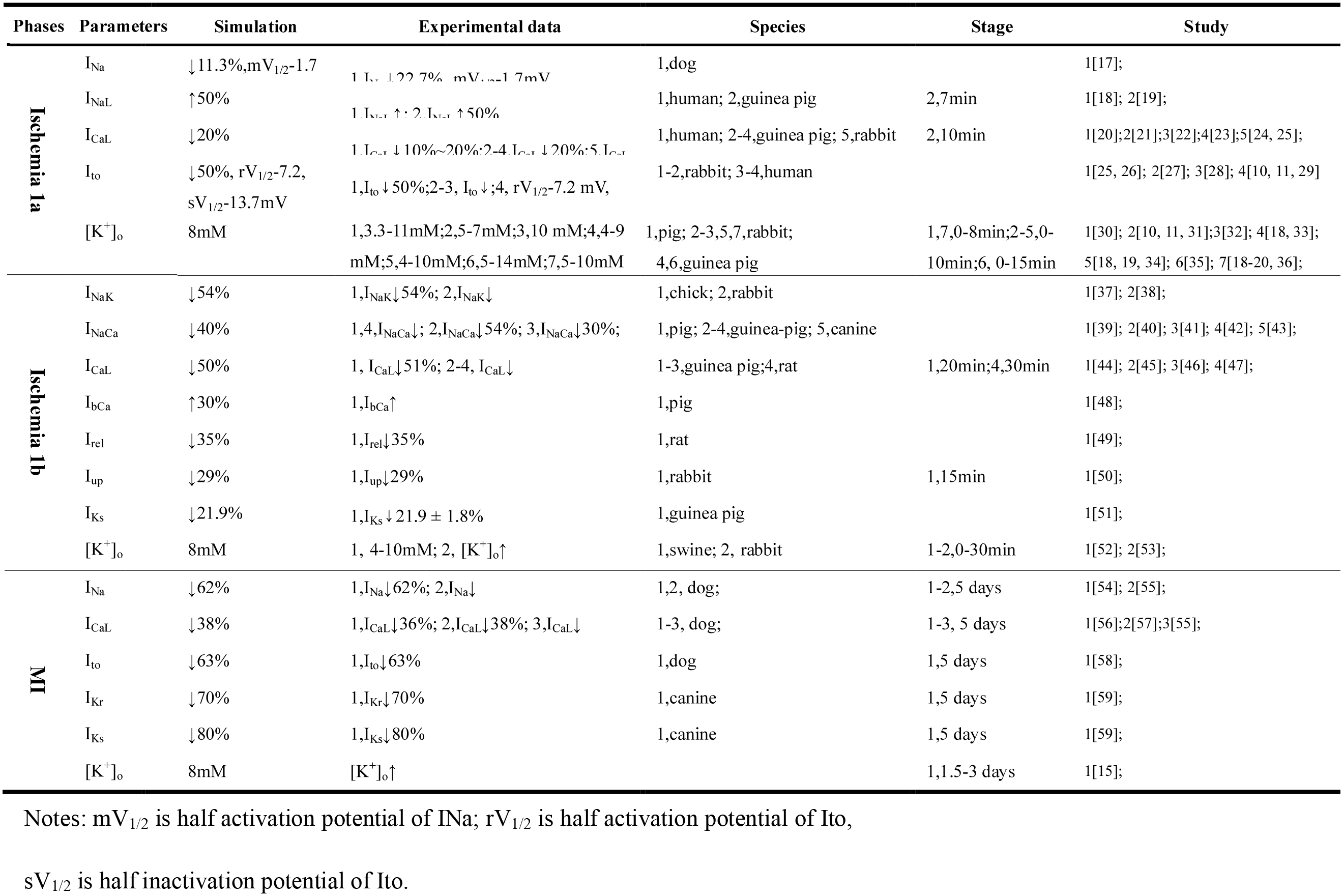
the electrophysiological properties of the three stages.

In addition to the ionic remodeling, the spatial heterogeneity was shown as a key factor of inducing reentry waves in myocardial ischemia or MI. In 2D regional ischemic tissues[4], Romero *et al.* studied effects of spatial heterogeneity distribution of ion currents and ion concentration on reentry wave propagation, and showed that the mismatch of the source–sink relationship was the ultimate cause of the unidirectional block leading to reentry, rather than the dispersion of refractoriness. Weiss *et al.* studied the effects of spatial heterogeneity distribution of endocardial (Endo), intermediate layer (M), and epicardial (Epi) cells on reentrant wave propagation, and linked features of the ECG to ionic mechanisms on ischemic tissues[5]. Several studies on MI revealed the effect of the spatial heterogeneity distribution of scar areas on the occurrence of reentry waves[8–11]. However, most of these studies focused on the heterogeneity induced by Endo, M, and Epi cells, and the impacts of spatial heterogeneity induced by differential remodeling on the reentrant waves remain unclear.

In order to better understanding the mechanisms of cardiac arrhythmias during evolution of coronary artery blockage, multi-scale computational models at the cellular, tissue, and organ levels based on the TP06 model were developed in this study. Simulation results in 2D tissues showed that the size of VWs was inversely proportional to the CV of the tissue, and the VWs in multiple pathological tissues were determined by the VWs in single pathological tissues. In addition, the VW of the tissue with three pathological conditions was larger than that in the tissue with ischemia 1a and 1b condition, and could be further enlarged by cell-to-cell decoupling. In order to reveal the mechanisms underlying the occurrence of reentry waves in multiple pathological tissues, two different spatial distributions of electrophysiological remodeling were constructed. In one case, the borders of different pathological areas are perpendicular to the excitation wavefront, and in the other case, the borders are parallel to the excitation wavefront. Reentrant waves were mainly induced by the spatial heterogeneity of refractory periods caused by the co-action of APD and CV along the wavefront when the spatial heterogeneity distribution is perpendicular to the excitation wavefront. When the borders were parallel to the wavefront, the elevated excitation threshold in MI region and the decline of the current transfer at gap junctions (I_gap_) were the primary reason for the generation of reentry. Finally, the simulation results on the 3D ventricle also demonstrated that the coexistence of multiple pathological conditions is more prone to induce reentry. The findings of this study sheds light on the arrhythmogenic mechanism during the progression of coronary artery blockage, and is helpful to develop new therapeutic strategies for arrhythmias in myocardial ischemia and infarction.

## Methods

### Single cell models

Based on the experimental data[12, 13], single cell models of ischemia 1a, 1b and MI were developed by incorporating the electrophysiological remodeling into the TP06 model. The experimental data of electrophysiological remodeling in different pathological conditions and the corresponding modifications of parameters in the computational models were summarized as shown in Table 1. In addition, in the simulations, the value of intracellular adenosine triphosphate concentration ([ATP]_i_) was 6.8 mM in normal condition and decreased to 4.6 mM in ischemia[14] and MI[15] stages according to the experimental data. The effect of [ATP]_i_ variation in pathological conditions on cell action potential was integrated into the model via I_KATP_ whose formula was proposed by Clayton *et al.*[16].

The stimulation current was applied on the single cell with a stimulation strength of −86.2 □pA/pF and a stimulus duration of 1□ ms. The time step was set as 0.02 □ms in our single cell simulation. A ventricular cell reached the steady state by 100 stimuli with a pacing duration of 1000 ms.

### 2D and 3D tissue models

The reaction-diffusion equation was used to model the mono-domain tissues[60]. The coupling between cells was isotropic in our simulations. The equation is modeled as follows:

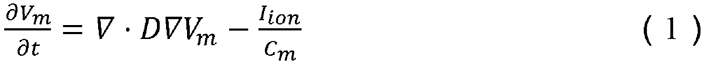

Where *V_m_* is the membrane potential, the value of the effective diffusion constant (D) is 0.154 *mm*^2^/*ms*, the value of the capacitance (*C_m_*) is 1*μF*/*cm*^2^,*I_ion_* is the total transmembrane current.

In addition to the remodeling in ionic currents, experimental data show that both ischemia1b[61–63] and MI[64] stages undergo a certain degree of cell-to-cell decoupling in the tissues (30∼40%↓), giving rise to a reduction in the CV of excitation wave in tissues[63]. The remodeling in cell-to-cell decoupling was mimicked by reducing the diffusion coefficient (30%↓) in ischemia 1b and MI stages.

The spatial distribution of various pathological condition in 2D and 3D tissues was shown in Fig 1. In 2D idealized tissue, the regions of pathological conditions along the direction of blood flow descendent are ischemia 1a, 1b, and MI regions, successively (Fig 1Ai and Aii). In order to investigate the generation of reentry waves in multiple pathological tissues, two different spatial distributions of electrophysiological remodeling were constructed. One distribution is shown in Fig 1Ai where ischemia 1a, 1b, and MI distributed horizontally and excitation waves propagated vertically. In this case, the boarders of various electrophysiological remodeling were perpendicular to the wavefront of excitation waves. On the contrary, as shown in Fig 1Aii, ischemia 1a, 1b, and MI distributed circularly in the other distribution. In this case when a stimulus was applied at the upper left corner, an excitation wave propagated circularly which is parallel to the boarders of various electrophysiological remodeling.

**Fig 1.**
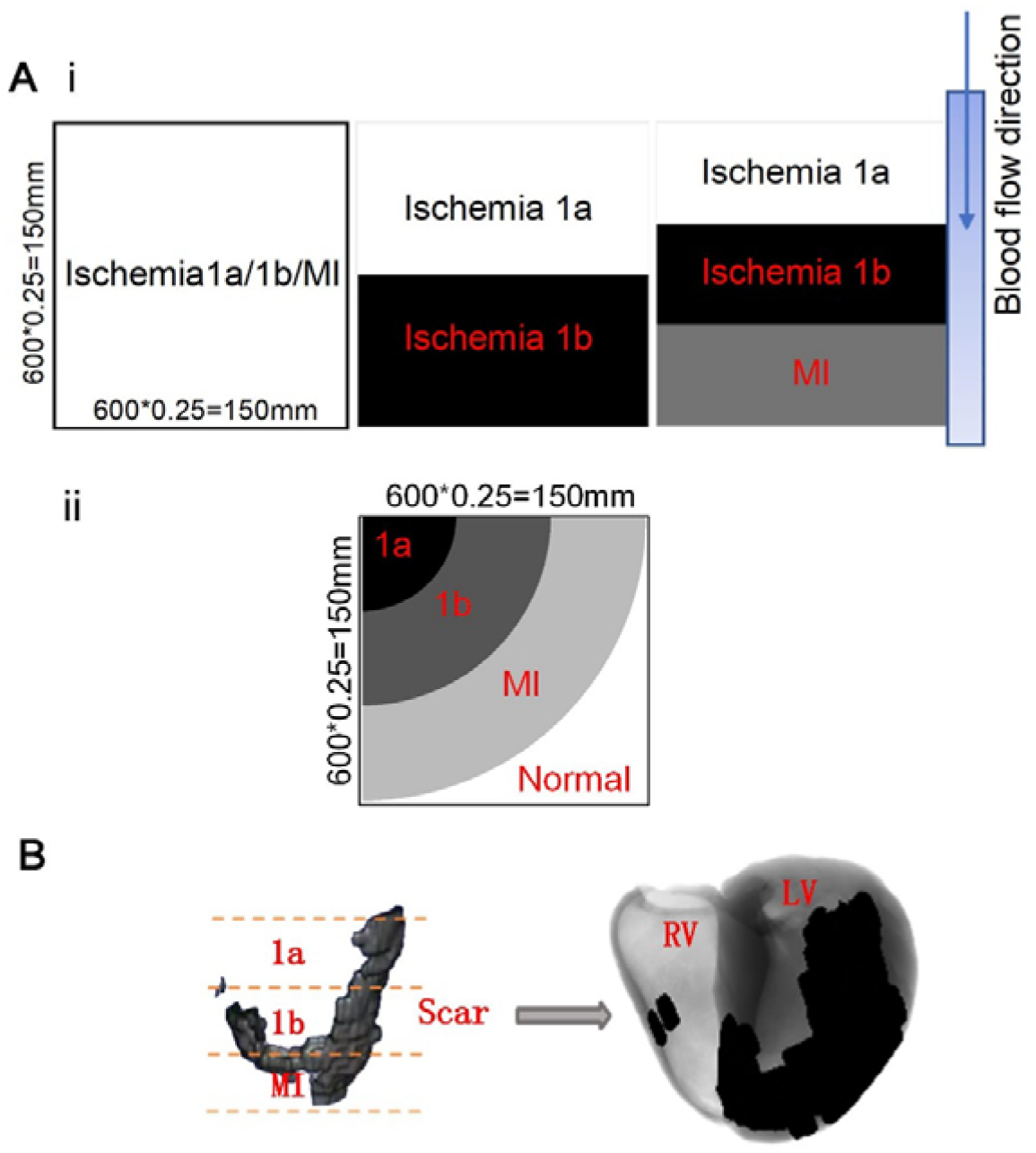
Tissue architecture of (A) 2D ideal tissues and (B) the 3D real ventricle tissue with ischemia 1a, 1b, MI and normal areas.

For 3D ventricular tissue model, 2D sections of the scar areas were extracted from the MRI images of a MI patient. There 2D slices were interpolated and reconstructed into a 3D scar which was then incorporated into the 3D human ventricle model according to our previous method[7]. Clinical studies have shown that obstruction of the anterior descending branch of the left coronary artery leads to ischemia of the anterior wall of the left ventricle and the interventricular septum[65, 66], which is consistent with the MI areas of this patient. Similar to the assumption of the vascular blood supply in the 2D tissue simulations, the blood flow direction of the coronary artery is from top to bottom. Therefore, the reconstructed scar in the 3D ventricular tissue were equally divided into three parts (from top to bottom: ischemia 1a, ischemia 1b, and MI) as shown in Fig 1B. The tissues outside the scar areas are normal tissues.

The calculation formula of safe factor (SF) improved by Romero *et al.* was used in this paper[67], as shown below:

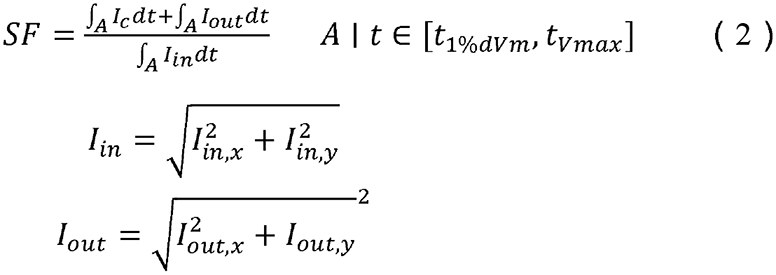

where I_in_ is the axial current entering the cell, I_out_ is the axial current leaving the cell, I_c_ is the capacitive current and A is the integration interval, from the instant when membrane potential derivative reaches 1% of its maximum (t_1%dVm_) and to the instant of membrane potential peak (t_Vmax_) during the depolarization phase.

### Stimulation protocols of 2D and 3D tissue models

The detailed information of stimulation protocols was shown in supporting information. The stimulation strength was −120 pA/pF in 2D and 3D simulations, and the stimulation duration was 3ms for 2D tissues and 2ms for 3D tissues. The 2D ideal tissues contained 600×600 nodes with a spatial resolution of 0.25 mm, resulting in a physical size of 150×150 mm^2^. In the 3D model, an isotropic domain composed of 325×325×425 nodes was used with a spatial resolution of 0.33 mm, corresponding to 107.25×107.25×140.25 mm^3^ in physical size. The standard S1-S2 protocol were used to initiate reentrant waves in the tissues and measure the VWs. For the S1-S2 protocol, five S1 stimuli with the interval of 1000 ms were applied on the leftmost three columns of nodes before S2 stimulus, which ensures the tissue reaching a steady state. S2 stimulus was applied in the upper left corner or lower left corner with the size of 300×300 cells. In the dynamic protocol, for the tissue in Fig 1Ai, stimuli were applied on the leftmost three columns of nodes, while, for the tissue in Fig 1Aii, the stimuli were applied on a circular sector at the upper left corner with a radius of 5 nodes. In the 3D ventricular tissue, stimuli were applied on a small cubic tissue with the size of 10×10×10 nodes in intramyocardial regions.

The differential equations were solved using the forward Euler method with a time step (Δt) of 0.02 ms. In 2D and 3D tissue models, the stability criterion and the Neumann boundary conditions were used[60]. An Intel core i7-3930K 64-bit CPU system was used to calculate the numerical solution, and parallel computing was applied for acceleration using Gtx titan z GPU.

## Results

### Electrophysiological remodeling in single cell models

The single cell models of different pathological conditions were validated via comparing the variations of electrophysiological characteristics with experimental data. Fig 2 showed that the variations of electrophysiological characteristics were similar in three pathological stages, such as the elevated resting potentials (RPs) (caused by hyperkalemia[30–34, 53]), the shortened APD (caused by the increase of I_KATP_[68, 69]), the decreased action potential amplitudes (APAs), and the reduced maximum depolarization rate (dV/dt_max_) (caused by the decrease of I_Na_[17]), implying an impaired excitability in these conditions. For ischemia 1a stage, the APD was 202.04ms which is within the range of human experimental data (180-260ms)[70]. For the ischemia phase 1b, the APD (146.86 ms) and APA (92.65 mV) were the smallest among these pathological stages, while the RP (−75.55 mV) was the highest. This may result in a spatial heterogeneity of the electrophysiological characteristics during the progression from stenosis to occlusion in the coronary artery. The APD in the ischemia 1b decreased by about half compared with that in the normal condition, which is basically consistent with the study of Pollard *et al.*[71]. In short, the simulation results of ischemia 1a and 1b are consistent with the previous studies[3, 16, 71, 72].

**Fig 2.**
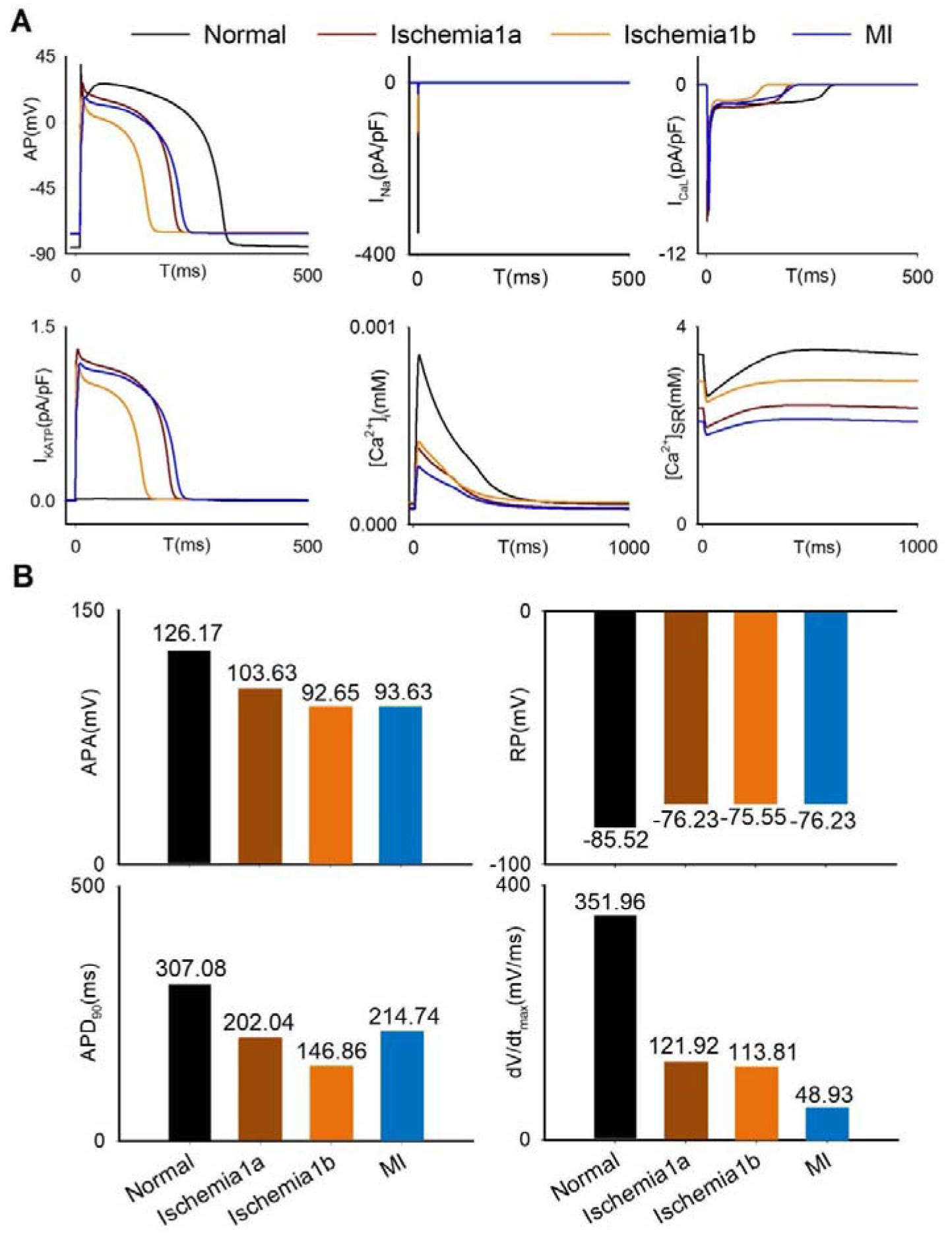
The variations of electrophysiological characteristics in (A) action potentials, several main ion currents, and (B) values of APA, RP, APD_90_ and dV/dt_max_ of single cells in normal, ischemia1a, ischemia1b, and MI.

For the MI stage, previous experimental studies have shown that the APD is shortened [58, 73–76]. However, it was shown that the APD was prolonged at the MI stage in many previous simulation studies[6, 10, 11, 77, 78]. Since it was reported that [ATP]_i_ was greatly decreased during the MI stage[15], in this study, this effect was incorporated into the model via I_KATP_ current activation induced by the decrease of [ATP]_i,_ leading to a shortened APD in the MI stage. As shown in Fig 2B, compared with the APD (307.08 ms) in normal situation, the APD (214.74 ms) in the MI phase decreased significantly, which is consistent with the experimental data[74]. Also, the APA (93.63 mV) and RP (−76.23 mV) in the MI phase were agree with experimental data[73]. Notably, compared with the normal condition, the dV/dt_max_ (48.93 mV/ms) decreased significantly in the MI stage and was the smallest among three pathological conditions as shown in Fig 2B. In summary, the simulation results of electrophysiology remodeling in three pathological conditions were consistent with the experimental data, which validated the developed single models of ischemia 1a, 1b, and MI stages.

### Vulnerability of 2D tissues to reentrant waves

VWs were measured in single and multiple pathological tissues to examine the effect of various remodeling on the vulnerability of cardiac tissue. In the single pathological tissues, the generation and maintenance of reentrant waves (S2 Fig) and the corresponding VWs (Fig 3B) were investigated using the S1-S2 cross-field protocol. The CV in each pathological condition decreased, comparing with that in normal conditions (Fig 3A), which is consistent with the relevant clinical trial records[63, 79]. Among the ischemia1a, 1b, and MI tissues, the CV in MI tissue was the lowest (31.25 cm/s), while the CV in ischemia 1b tissue was the highest (52.25 cm/s) as shown in Fig 3A. In addition, inevitably, cell-to-cell decoupling further reduced the CVs of the tissue with ischemia 1b and MI to 42.75 cm/s and 25.5 cm/s, respectively. Meanwhile, compared with the VW of normal tissues (110 ms), the VWs in tissues with ischemia 1a, 1b, and MI alone increased to 150 ms, 140 ms, and 230 ms, respectively. Cell-to-cell decoupling further enlarged the VWs of the tissue with ischemia 1b and MI to 170 ms and 280 ms, respectively. These simulation results showed that the size of VWs is inversely proportional to the CV of the tissue, which is consistent with previous study [80].

**Fig 3.**
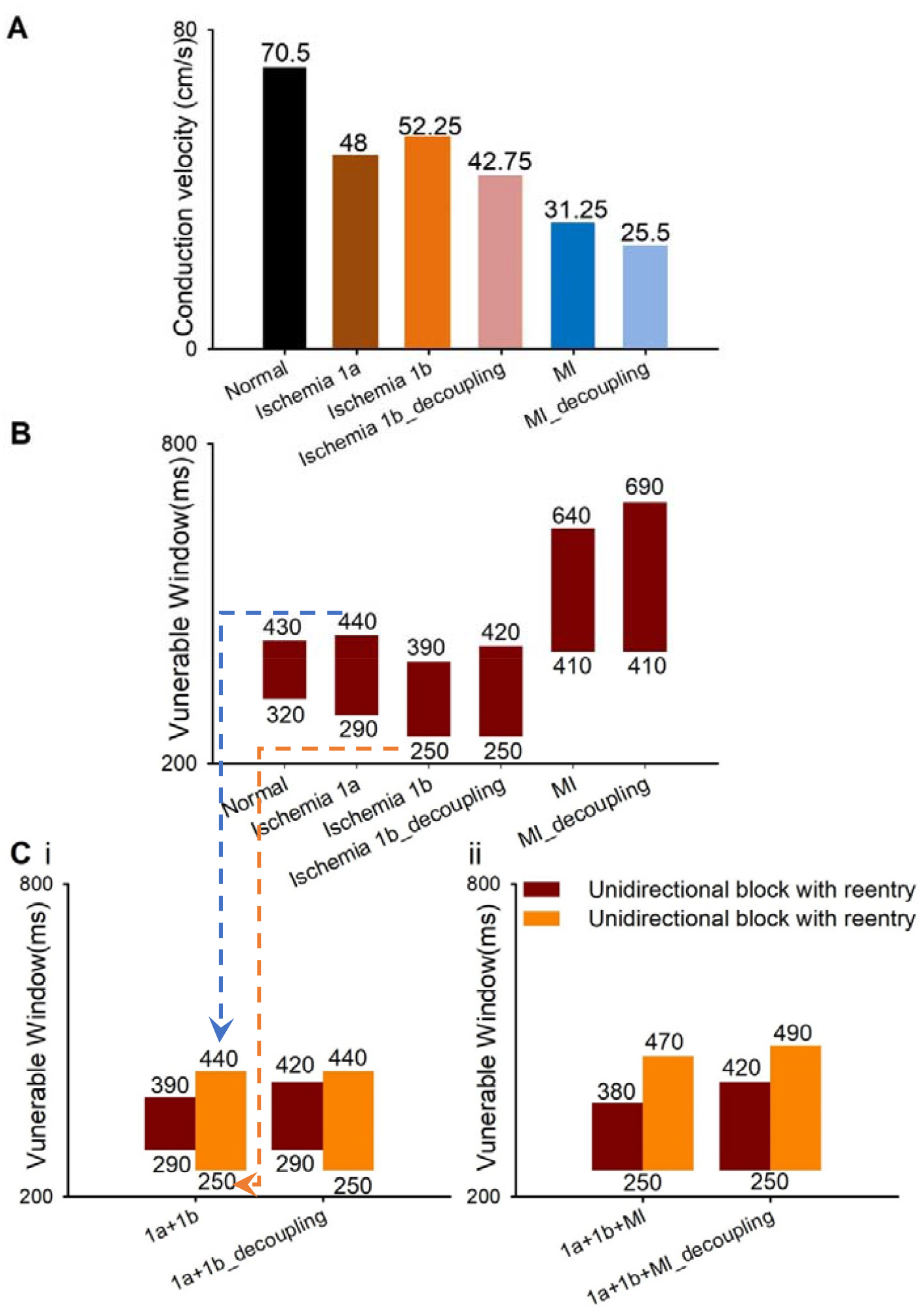
(A) The CV in six different 2D homogenous tissues: normal, ischemia 1a, 1b, MI, decoupled 1b and decoupled MI. (B) The VWs of 2D homogenous tissues when the S2 stimulus was applied in the lower left corner using the S1-S2 protocol. (C) The VWs of 2D multiple pathological tissues (i) where ischemia 1a and 1b distributed horizontally and (ii) where ischemia 1a, 1b, and MI distributed horizontally when the S2 stimulus was applied in the (Dark Red) lower left or (Orange) upper left corner using the S1-S2 protocol.

Then, the reentrant waves and VWs were investigated using the S1-S2 cross-field protocol in the inhomogeneous tissues with multiple pathological conditions. In the tissue with two pathological conditions (ischemia1a and 1b/decoupled 1b), the S2 stimulus was applied in the upper or lower left corner, which is corresponding to ischemia 1a or 1b region, respectively (Fig 1Ai). When the S2 stimulus was applied in ischemia 1a regions (upper left corner), the upper bound of the VW of the multiple pathological tissue (440 ms) was the upper bound of the VW of the homogenous ischemia 1a tissue. Meanwhile the lower bound of the VW of the multiple pathological tissue (250 ms) was the lower bound of the VW of the homogenous ischemia 1b tissue (Fig 3Ci, orange bars). Intriguingly, on the contrary, when the stimulus was applied in ischemia 1b regions (lower left corner), the upper and lower bound of the VW of the inhomogeneous tissue was the upper bound of ischemia 1b tissue and the lower bound of the ischemia 1a tissue, respectively (Fig 3Ci, red bars). This hypothesis still held true when cell-to-cell decoupling occurred in the ischemia 1b region (Fig 3Ci). Therefore, the VW of multiple pathologies tissues is determined by the upper and lower bounds of VWs of single pathological tissues.

In the above hypothesis, when the S2 stimulus was applied in ischemia 1b regions (lower left corner, Fig 4A), the lower limit of the VW of multiple pathological tissue is the limit allowing upward unidirectional conduction, i.e. the lower limit of the VW of the homogenous ischemia 1a tissue (Fig 4A, pacing cycle length (PCL)=290ms). The upper limit of the VW of multiple pathological tissue is the limit allowing bidirectional conduction of S2 stimulus, i.e. the upper limit of the VW of the homogenous ischemia 1b tissue (Fig 4A, PCL=420ms). Similarly, when the S2 stimulus was applied in ischemia 1a regions (upper left corner, Fig 4B), the VW is determined by the limits of downward unidirectional conduction (Fig 4B, PCL=250ms) and bidirectional conduction (Fig 4B, PCL=440ms) of S2 stimulus. In the tissue with three pathological conditions (ischemia1a, 1b, and MI), the VWs (Fig 3Cii) were not simplify determined by the above conclusion. In this case, the lower limit of the VW of the multiple pathological tissue is the limit allowing upward/downwards unidirectional conduction, i.e. the lower limit of the VW of the homogenous ischemia 1b tissue (Fig 3Cii, 250ms). However, the limit allowing bidirectional conduction of S2 stimulus (the upper limit of the VW) is influenced by the excitation wave propagation across two regions (ischemia 1a and 1b or ischemia 1b and MI, S3 Fig). Although the upper limit of the tissue with three pathological conditions could not be precisely determined, the VW of the tissue with three pathological conditions was larger than that in the tissue with ischemia 1a and 1b condition (Fig 3Ci and Cii).

**Fig 4.**
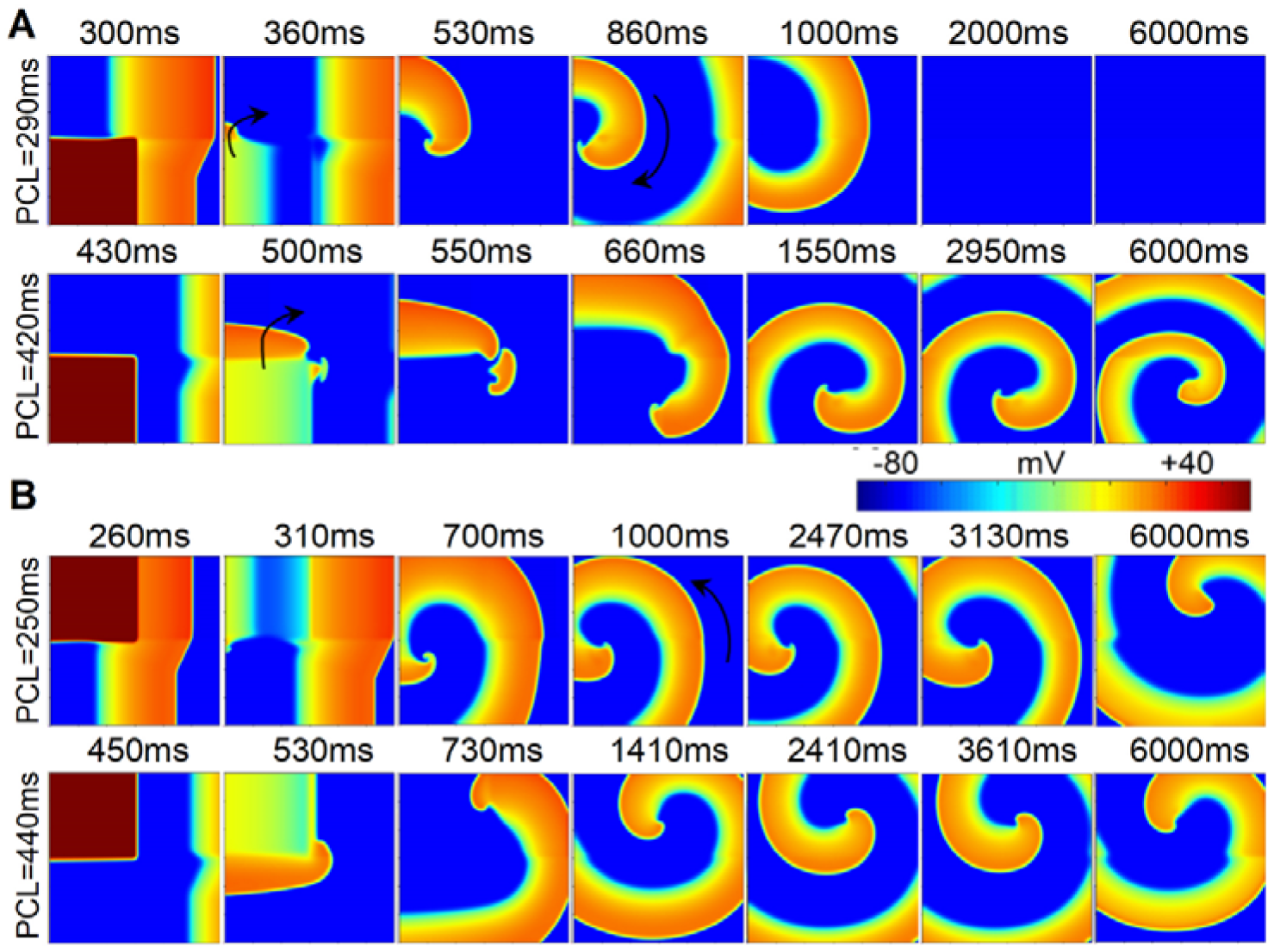
Wave propagation in the 2D tissue where ischemia 1a and decoupled ischemia 1b distributed horizontally using the S1-S2 protocol when the S2 stimulus was applied in the (A) lower left or (B) upper left corner.

### Initiation of reentrant waves in 2D multiple pathological tissues

In order to reveal the mechanisms underlying the genesis of reentry waves in 2D tissues, excitation wave propagation was investigated in multiple pathological tissues with two different distributions. Firstly, the reentrant waves were investigated when the spatial heterogeneity distribution was perpendicular to the excitation wavefront. In this section, both S1 and S2 stimuli were applied to the leftmost 3-line cells to induce reentrant waves (Fig 5). Different from the previous section, using this stimulation mode, reentrant waves were induced in multiple pathological tissues with ischemia 1a, 1b, and MI, but not in homogeneous 2D tissues. The VWs of occurrence reentrant waves were explored using the S1-S2 protocol (Fig 5A). In this case, the reentrant wave was induced at the border between ischemia 1b and MI regions due to the conduction block of S2 in MI area. The unidirectional conduction was accounted for the long refractory period of MI induced by the low CV in MI region (Fig 5Ai). Surprisingly, the size of VWs was the same in coupling and decoupling conditions (as shown in Fig 5Aii). This may be explained by the fact that in the cell-to-cell decoupling condition, although the CVs decreased in both ischemia 1b and MI regions, the variation of CV across their border remained unchanged, i.e., the decline in CV across the border was ∼40% in coupling (52.25 to 31.25 cm/s, Fig 3A) and decoupling (42.75 to 25.5 cm/s, Fig 3A) conditions. When stimuli were applied to the leftmost 3-line cells using the dynamic stimulation protocol with a pacing cycle of 250ms (the detail protocol was shown in supplemental materials), spiral wave breakup occurred in the 2D tissue (Fig 5Bi), leading to severely deformed action potentials at point P2 (Fig 5Bii) and thus an arrhythmic pseudo-ECG of the tissue (Fig 5Biii).

**Fig 5.**
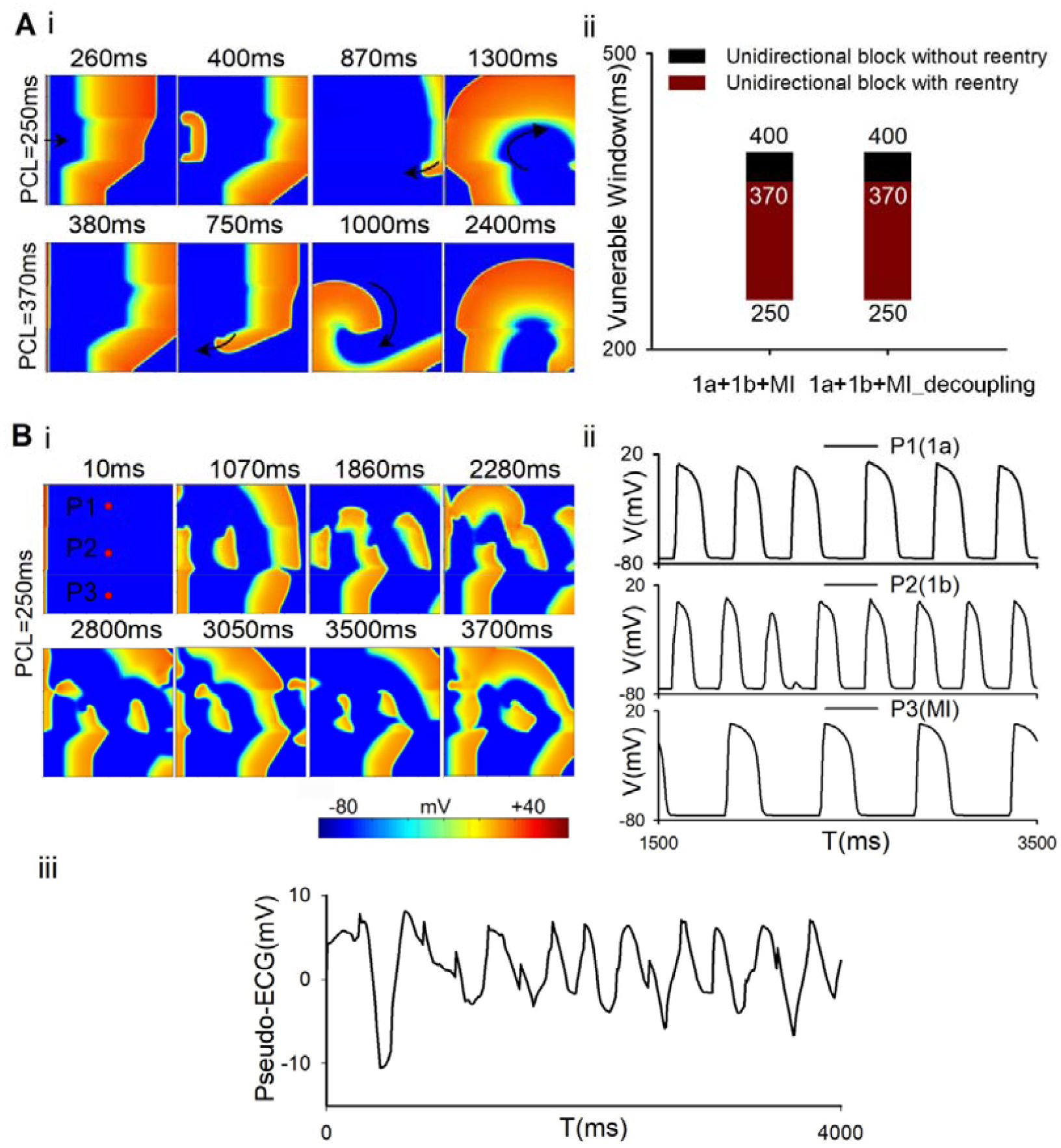
(A) (i) Wave propagation in the 2D tissue where ischemia 1a, decoupled 1b, and decoupled MI distributed horizontally and (ii) corresponding VWs when the leftmost stimulation was applied using S1-S2 protocol. (B) (i) Wave propagation, (ii) action potentials of points P1, P2 and P3 and (iii) Pseudo-ECG in the 2D tissue when the leftmost stimulation was applied using dynamic stimulus protocol.

Fig 6A showed the spatial distribution of APD in the 2D tissue was perpendicular to the excitation wavefront which indicated by the line L1 in Fig 6Ai. It was shown that the largest APD gradient is located at the junction of different pathological tissues (Fig 6Aii). The difference of APD between ischemia 1a and 1b was similar to that between ischemia 1b and MI (Fig 6B), but the decline in CV between ischemia 1b and MI was much greater than that between ischemia 1a and 1b (Fig 6C). It suggested that multiple electrophysiological remodeling of various pathological conditions produced sharp heterogeneity along the excitation wavefront (the line L1). The formation of reentry waves is mainly due to the difference of CV between ischemia 1b and MI (see the details in Discussion).

**Fig 6.**
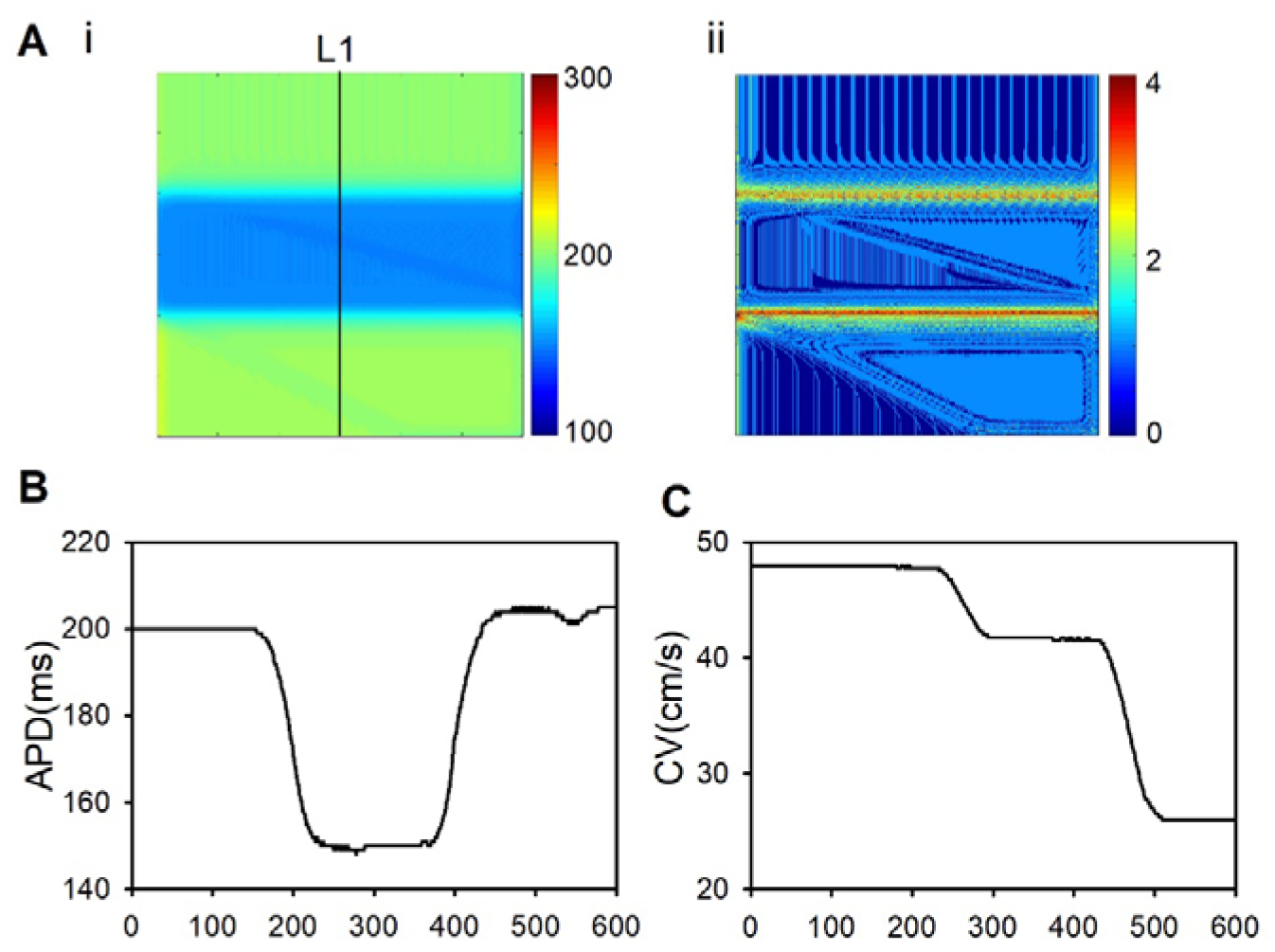
(A) (i) APD distribution of the first stimulation in the 2D tissue where ischemia 1a, decoupled 1b, and decoupled MI distributed horizontally when the leftmost stimulation was applied with a pacing cycle of 250ms. (ii) The maximum APD difference between each cell and its neighbors in the 2D tissue. (B) APD of all cells along the line L1 in the 2D tissue. (C) CV of all cells along the line L1.

In the other spatial heterogeneity distribution, the borders of different pathological regions were parallel to the excitation wavefront (as shown in Fig 7A). In this case, excitation wave propagation in the 2D multiple pathological tissues was investigated using the dynamic stimulation protocol. Fig 7Bi showed the development of reentrant waves in the 2D tissue with ischemia 1a, decoupled 1b, and decoupled MI when the stimulation interval was 420ms. In this process, the excitation wave was blocked at marginal areas in MI region (Fig 7Bi, 2630 ms). With the propagation of the excitation waves along the diagonal line, the excitation site of the 6th stimulus located on the boundary between MI and normal areas (Fig 7Bi, 2800 ms) resulted in backward waves (as shown by the black arrows in Fig 7Bi, 2920 ms) and generated the reentrant waves. Then, the reentrant wave collided with the excitation wave of the 7th stimulus, hindering the continued wave propagation. This process repeated in the following stimuli (Fig 7Bi, 3350-3600ms).

**Fig 7.**
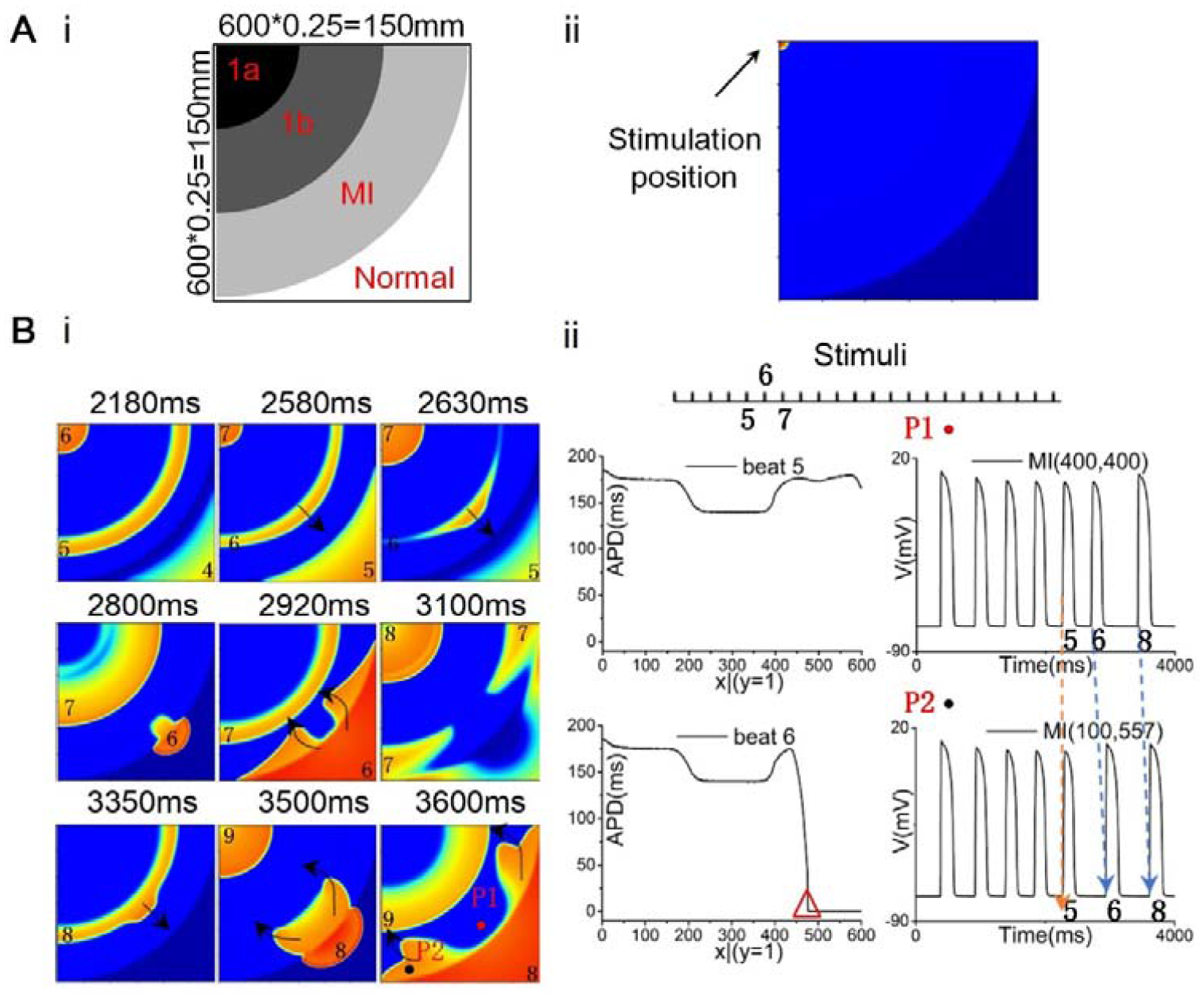
(A) (i) The 2D tissue structure where ischemia 1a, ischemia 1b and MI distributed circularly and (ii) the location of point stimulation. (B) (i) Wave propagation in the 2D tissue with ischemia 1a, decoupled 1b, and decoupled MI. (ii) APD of all cells along the border of the 2D tissue at the 5th and 6th stimulation and action potentials at points P1 and P2.

Different from the sharp heterogeneity along the excitation wavefront when the spatial heterogeneity distribution was perpendicular to the excitation wavefront (Fig 6), S4 Fig showed the spatial heterogeneity of APD along the excitation wavefront was trivial when the spatial heterogeneity distribution was parallel to the excitation wavefront. In order to reveal the mechanisms of the genesis of reentrant waves in this condition, the distribution of SF in the tissue was calculated. As shown in Fig 8, the SFs along Arc1(Fig 8A) were slightly smaller at the edge of the tissue (0° and 90°) in the 5th stimulation. This small variation in SF along the excitation wavefront gradually diminished (as shown in Arc 2, Fig 8A), and the excitation propagated normally in the 5th stimulation. However, in the 6th stimulation, the SFs at the edge of the tissue along Arc1(Fig 8B) further decreased to 0.875 (<1) and, therefore, led to blockage of excitation waves at the edge (as shown in Fig 7Bi, 2630ms), which contributed to the backward excitation waves (Fig 8B) and induced reentry. The declined SFs at the edge areas mainly resulted from the decreased I_gap_ at the edge area along the excitation wavefront (Fig 8Cii).

**Fig 8.**
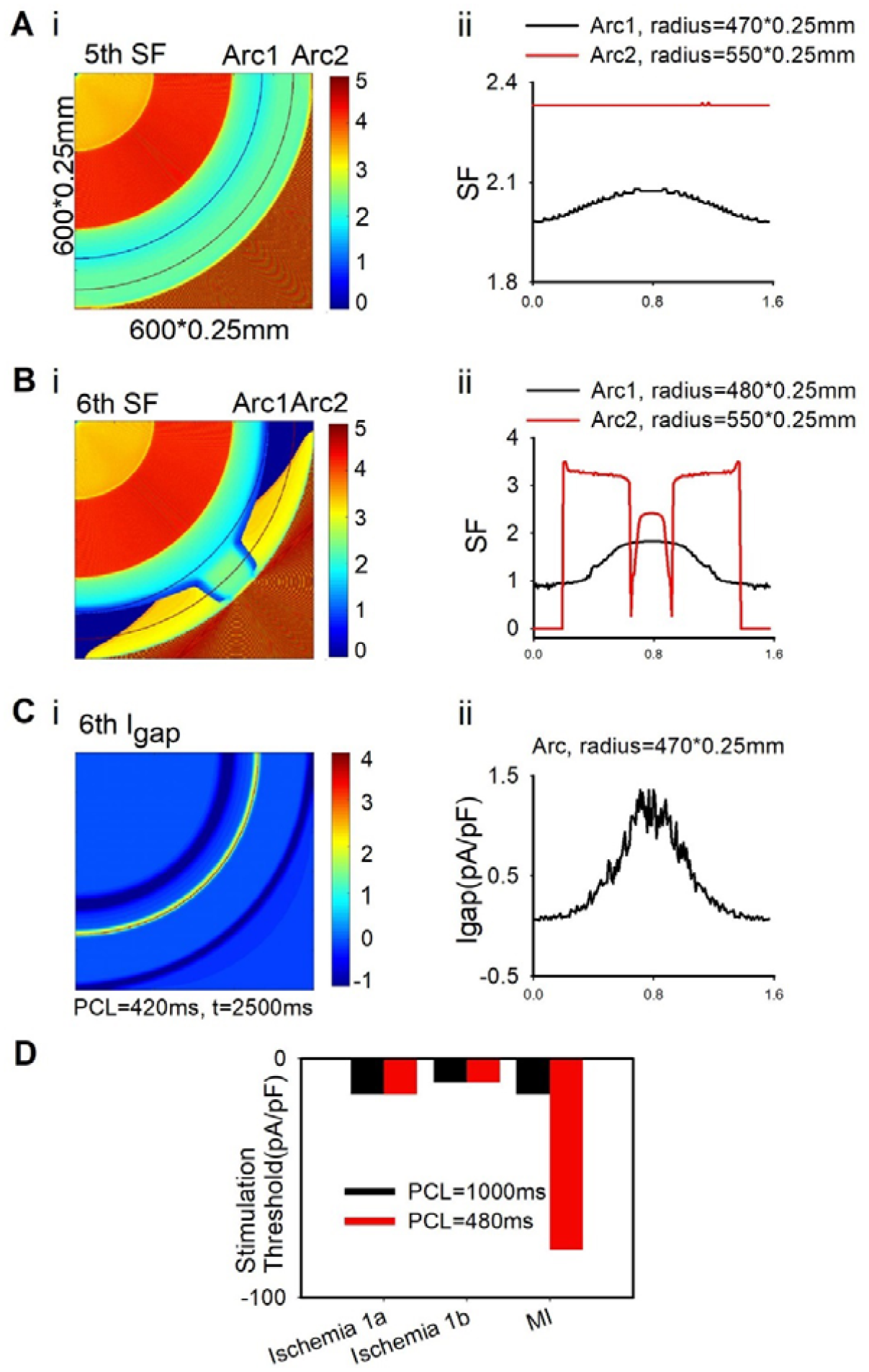
The SF distribution in the 2D tissue where ischemia 1a, decoupled 1b, and decoupled MI distributed circularly at the (Ai) 5th and (Bi) 6th stimulation and (Aii and Bii) the SFs of all cells along Arc1 and Arc2. (C) (i) Igap distribution in the 2D tissue and (ii) Igap of all cells along the arc at 6th stimulation. (D) The minimum stimulation current for tissue excitability in the 2D tissue at different PCL (480ms; 1000ms) in ischemia 1a, 1b and MI conditions with point stimulation in the upper left corner.

### Reentrant waves in 3D tissue model

Fig 9A showed that in normal conditions, reentrant waves did not occur after S1, S2 and S3 stimulation. However, in 3D ventricular tissues with multiple pathological conditions (Fig 1B), the slowed CVs in the pathological regions resulted in a slowed and heterogeneous repolarization (as shown in Fig 9B, 560ms). When the S3 stimulus was applied, the unidirectional blockage of wave propagation occurred (as shown in Fig 9B, 600ms), which in turn triggered reentrant waves (Fig 9B, 900ms). When the reentrant waves arrived the scar areas again, wavefront breakup was induced, giving rise to more disordered reentrant waves (Fig 9B, 1800ms). The reentrant waves persistently existed within 26000ms. Notably, using the same stimulation mode, reentrant waves were induced in the tissue with ischemia 1a only, but the duration of existence was relatively short (8000 ms) as shown in S5 Fig. Moreover, no reentry wave was induced in the tissue with ischemia 1b or MI only as shown in S5 Fig. It suggested that the coexistence of multiple pathological conditions in the tissue was more prone to cardiac arrhythmias.

**Fig 9.**
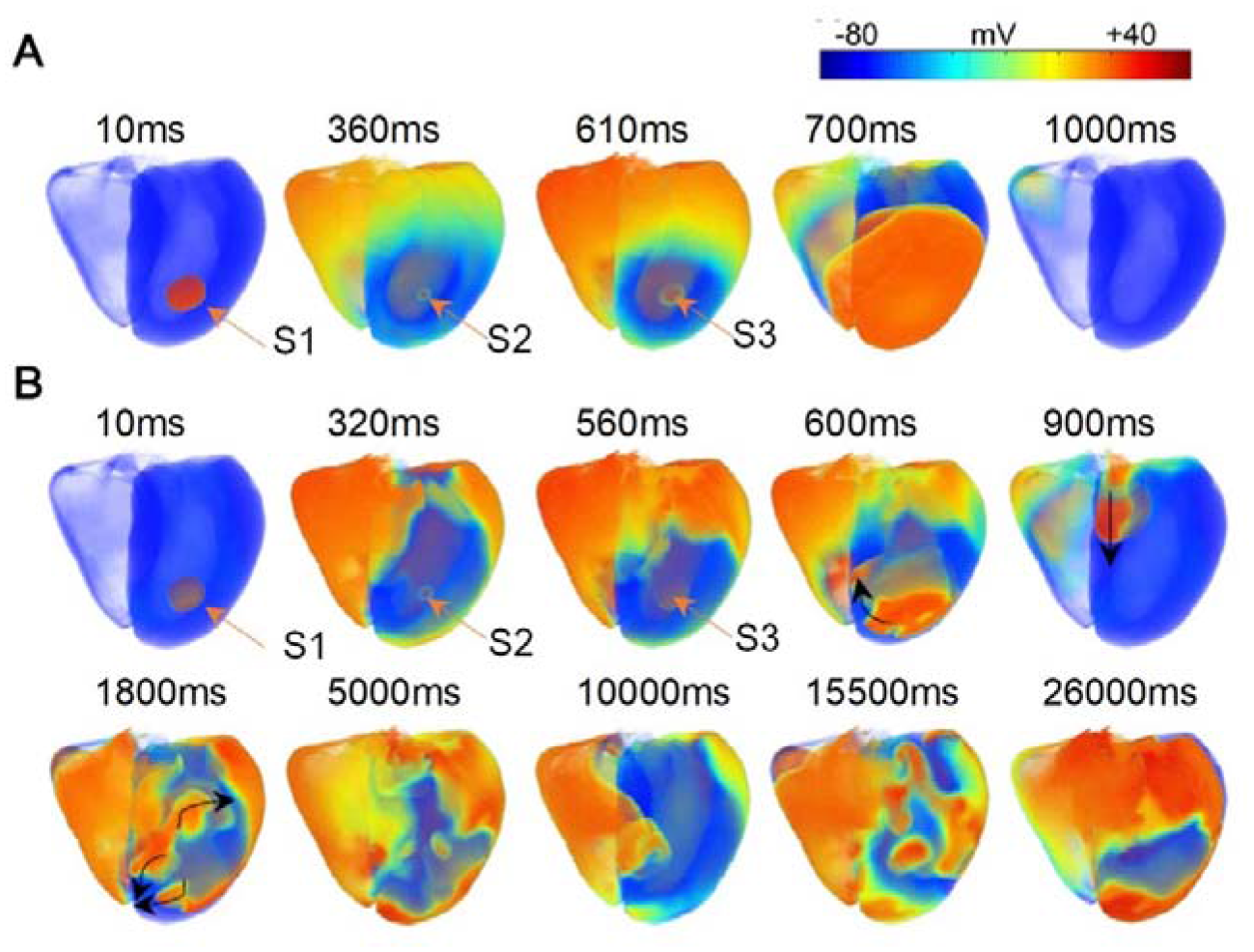
(A) Wave propagation in the 3D ventricular tissue in normal conditions (with stimulus interval 350ms and 250ms, respectively). (B) Wave propagation in the 3D ventricular tissue with the ischemia 1a, 1b, and MI (with stimulus interval 310ms and 240ms, respectively).

## Discussion

With the development of ischemia, different electrophysiological remodeling occurred over time. Many studies have explored the effects of electrophysiological remodeling and the heterogeneous distribution of tissues on the wave propagation in the ischemia [4, 81] and infarcted tissues[5, 9]. However, the mechanisms of initiation of reentrant waves in the tissue with multiple ischemic remodeling are not fully understood. In this paper, the mechanism of cardiac arrhythmias caused by myocardial ischemia and MI during coronary artery occlusion was systematically investigated. The main findings of this studies are: (1) Computational models of ischemia 1a, 1b, and MI stages were developed and validated using experimental data. (2) The size of VWs was inversely proportional to the CV of the tissue. In 2D tissue, the VW of multiple pathological tissue was determined by the VWs of single pathologic tissues. In addition, in the tissue with three pathological conditions, the VW was larger than that in the tissue with ischemia 1a and 1b condition, and could be further enlarged by cell-to-cell decoupling. (3) When the spatial heterogeneity distribution is perpendicular to the excitation wavefront, the formation of reentrant waves is caused by the spatial heterogeneity of refractory periods due to the interaction of APD and CV along the wavefront. And, the heterogeneity of the CV between ischemia 1b and MI on the wavefront plays a major role. (4) When the spatial heterogeneity distribution is parallel to the excitation wavefront, the elevated excitation threshold in MI region resulted in the decline of I_gap_ and the conduction block in edge of the tissue along the excitation wavefront, producing the reentrant waves.

### The properties of the VW of multiple pathological tissues

In multiple pathological tissue, the VW, measured using the S1-S2 cross-field protocol, was different when the S2 stimulus was applied at lower or upper left corner (Fig 3C). The size of VWs was determined by the boundaries of individual VWs in single pathological tissue. For the S1-S2 cross-field protocol, the upper and lower boundaries of the VW were determined by the time limits of bidirectional (horizontal) and unidirectional (vertical) propagation of S2 stimulus, respectively. In the tissue with two pathological conditions (i.e. ischemia 1a and 1b, Fig 4), S2 stimulus located in ischemia 1a (upper left corner) or 1b (lower left corner) region. Assuming S2 stimulus was applied in 1b (lower left corner) region, the time limits of horizontal and vertical propagation of S2 stimulus were the upper boundary of the VW in ischemia 1b tissue and the lower boundary of the VW in ischemia 1a tissue (Fig 3Ci, orange bar). Similar conclusion can be derived when S2 stimulus was applied in ischemia 1a (upper left corner) region. In the tissue with three pathological conditions (ischemia 1a, 1b, and MI), the lower boundary of the VW of multiple pathological tissue is the limit allowing vertically unidirectional conduction of S2 stimulus (upwards/downwards, S3 Fig), i.e. the lower limit of the VW of the homogenous ischemia 1b tissue (Fig 3Cii). However, the limit allowing bidirectional conduction of S2 stimulus (the upper limit of the VW) was the limit allowing horizontal propagation of S2 stimulus. In this case, the horizontal propagation of S2 stimulus was influenced by the two pathological regions (ischemia 1a and 1b or ischemia 1b and MI, S3 Fig), and, therefore, cannot be derived precisely. In addition, another conclusion was that the VWs were enlarged by cell-to-cell decoupling in the tissues with two or three pathological conditions (Fig 3C), and the VW of the tissue with three pathological conditions was larger than that in the tissue with ischemia 1a and 1b condition (Fig 3C), suggesting a higher risk of arrhythmias.

### The mechanism of reentry initiation when the spatial heterogeneity distribution is perpendicular to the excitation wave front

When the stimulation was applied to the leftmost side of the tissue, there was no reentry wave in the single pathological tissue with ischemia 1a, 1b, or MI alone. Also, reentry was not observed in the tissue with both ischemia 1a and 1b, but was seen in the tissue with ischemia 1a, 1b, and MI (as shown in Fig 5). The VW declined in the tissue with ischemia 1a, 1b, and MI, when ischemia 1b area was replaced with ischemia 1a area (as shown in S6 Fig). When the spatial heterogeneity distribution is perpendicular to the excitation wavefront, sharp variations in electrophysiological characteristics (APD and CV) were produced at the border between different pathological conditions along excitation wavefront (Fig 6). As the excitation wave length (WL) is calculated as the product of APD and CV, the excitability and refractoriness of the tissue at the border between different pathological conditions were altered significantly. Due to the decline in APD and CV, the WL was shortened in ischemia 1b compared to that in ischemia 1a region (Fig 5Ai, 260ms). Meanwhile, the declined CV resulted in a slowed wave propagation. These two effects counterbalanced with each other, leading to a relatively little time difference between wave tails in ischemia 1a and 1b region (Fig 5Ai, 260ms). At the border between ischemia 1b and MI, although the WL was further shortened in MI region (Fig 5Ai, 260ms), the wave tail of MI region was significantly delayed (Fig 5Ai, 260ms) due to the sharp decline in CV (Fig 6C). The delayed wave propagation increased the refractory period in MI region, and was responsible for the initiation of reentrant waves (Fig 5). Therefore, the heterogeneity along the excitation wavefront was the main reason of reentry, when the spatial heterogeneity distribution is perpendicular to the excitation wavefront. The heterogeneity of the CV on the wavefront plays an important role. Especially, the relative difference of the CV between ischemia 1b and MI is the main determinant.

### Initiation of reentrant waves in the tissue with gradient heterogeneity

In the previous section, the region-wise heterogeneity caused sharp variations in electrophysiological characteristics at the borders, which was rarely seen in real heart. Therefore, it was assumed that the variations in electrophysiological characteristics across the tissue was gradually induced with the progress of myocardial ischemia, implicating a gradient-wise heterogeneity. Reentrant waves were observed when I_KATP_ and I_Na_ were set as gradient distribution individually as shown in Fig 10 (see details in supporting information). As I_KATP_ and I_Na_ are closely associated with the APD and CV, respectively, the gradient-wise heterogeneity of I_KATP_ or I_Na_ corresponded to a decreased heterogeneity of APD or CV along the wavefront, respectively. As shown in Fig 10B, although both effects led to a reduced VW, a gradient distribution of I_Na_ had a more profound effect in reducing the VW. This suggested that the heterogeneity of the CV, rather than the APD, was the main factor of the genesis of reentry, which was consistent with the conclusion in region-wise heterogeneity. Although reentry was not induced when all the remodeled parameters were set as gradient distribution, breakup of reentry waves was induced using the dynamic stimulation protocol (as shown in S7 Fig), indicating the potential risk of arrhythmia.

**Fig 10.**
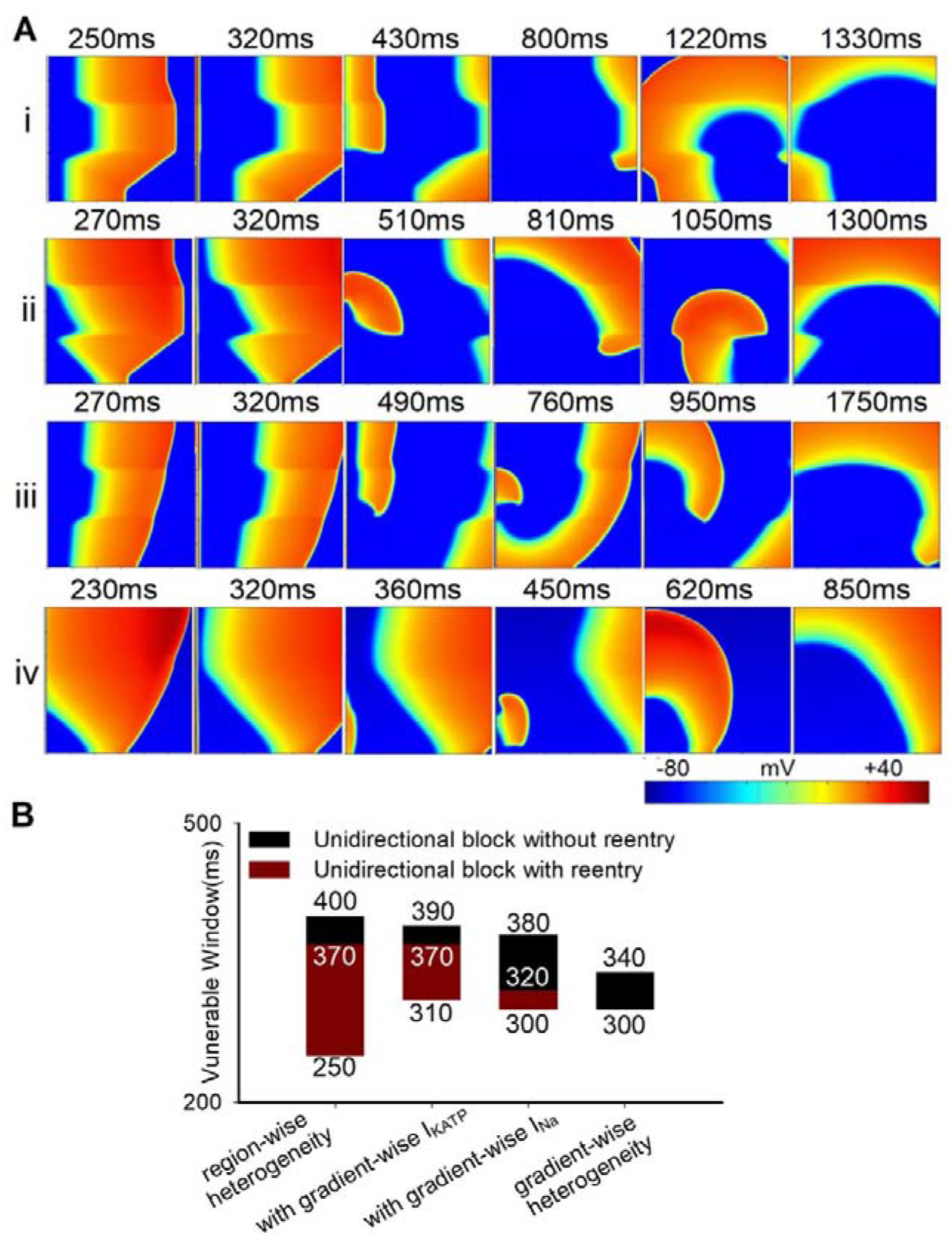
(A) Wave propagation in the 2D tissue with ischemia 1a, 1b, and MI and (B) VWs when the leftmost stimulation was applied using S1-S2 protocol in four different conditions: (i) region-wise heterogeneity, (ii) with gradient-wise I_KATP_, (iii) with gradient-wise I_Na_, (iv) gradient-wise heterogeneity of all parameters.

### The mechanism of reentry initiation when the spatial heterogeneity distribution is parallel to the excitation wavefront

When the spatial heterogeneity distribution is parallel to the excitation wavefront, there was no spatial heterogeneity along the excitation wavefront at the beginning of stimulation. The simulation results showed that I_gap_ was smaller at the edge of the tissue (Fig 8C), which may be due to less cells were coupled at the edge, leading to a weaker drive force for excitation. Moreover, it was shown that the excitation threshold in MI region was dramatically increased at high pacing frequency (Fig 8D), implicating that the drive force required for excitation in MI region was enlarged. In this case, with the fast stimulation using the dynamic stimulation protocol, the safe factor for conduction at the edge in MI region was gradually decreased (Fig 8A and 8B), which eventually gave rise to the conduction block at the edge and promote the formation of reentrant waves (Fig 7).

What’s more, if the ischemia 1b area was replaced with the ischemia 1a area, the VW declined from 50 ms (420 ms∼470 ms) to 40 ms (430 ms∼470 ms), when the spatial heterogeneity distribution is parallel to the excitation wavefront. As shown in Figs 2 and 3, compared with ischemia 1a stage, the lower APA, dV/dt_max_ and CV as well as the higher RP impaired the excitability of ischemia 1b region, impeded the wave propagation. Therefore, even the electrophysiological remodeling is homogeneous along the excitation wavefront, the axial spatial heterogeneity caused by multiple pathological conditions may hinder the wave propagation, further reduced the drive force for excitation, and contributed to the conduction block at the edge in MI region. As the edge of ischemic region in real heart tissue is much more complicated than the ideal tissue in this study, the risk of reentrant arrhythmia would be higher when multiple pathological conditions coexisted.

### Reentrant waves in 3D tissue

Reentrant arrhythmias occurring on the ventricles are mainly ventricular tachycardia and ventricular fibrillation, and ventricular fibrillation is one of the most dangerous arrhythmias, the hallmark of which is the appearance of wave fissures[82]. In the 3D ventricular tissue, persistent reentrant waves were observed in the tissue with three pathological conditions, rather than in the single pathological tissue with ischemia 1a, 1b, or MI. This phenomenon was consistent with our conclusion in the previous section that in multiple pathological tissue using the leftmost stimulation, the enhanced spatial heterogeneity of the tissue promoted the generation of reentry waves, thus greatly increased the risk of arrhythmia. In addition, the anatomical heterogeneity in real heart tissue further amplified the risk of reentrant arrhythmia in multiple pathological conditions, since it would augment the spatial heterogeneity along the wavefront as well as along the edge of ischemic region.

## Limitation

The limitations of this paper exist in the following aspects. At single cell-level, the limitations of TP06 cell model also exist in our simulation, for example, the effect of CaMKII is not included. In addition, it depends on the further development of patch-clamp experimental studies of human cells in ischemia 1a, 1b, and MI, so that more real human data can be used to remodel single cells. In addition, the effect of the heterogeneity of the three cardiomyocyte cells (Endo, M, Epi cells) was not compared in our single-cell and 2D simulation experiments, only the Epi cells were used. However, this does not change the conclusions in single-cell and 2D tissue simulations, and these three different cells were included in our 3D real ventricular tissues. The stage of MI is often accompanied by fibrosis, while fibrosis tissues were not added into our models, which can be supplemented in the subsequent simulation work. Furthermore, more diversified and complex distribution of ischemia 1a, 1b, and MI areas may be designed in 2D tissues and 3D tissues, but this will not change our conclusions.

## Author Contributions

Conceptualization: Qince Li, Henggui Zhang.

Data curation: Cuiping Liang, Qince Li.

Formal analysis: Cuiping Liang, Qince Li, Kuanquan Wang, Henggui Zhang.

Funding acquisition: Yimei Du, Qince Li, Kuanquan Wang.

Investigation: Cuiping Liang, Qince Li, Kuanquan Wang.

Methodology: Cuiping Liang, Qince Li, Henggui Zhang.

Project administration: Qince Li, Kuanquan Wang, Henggui Zhang.

Resources: Qince Li, Kuanquan Wang, Henggui Zhang.

Software: Cuiping Liang, Henggui Zhang.

Supervision: Qince Li, Kuanquan Wang, Henggui Zhang.

Writing – original draft: Cuiping Liang, Qince Li.

Writing – review & editing: Cuiping Liang, Qince Li, Kuanquan Wang, Yimei Du,

Wei Wang, Henggui Zhang.

## Supporting information

**S1 Fig.** The variations of electrophysiological characteristics of other ion currents and concentration of single cells in normal, ischemia1a, ischemia1b, and MI conditions.

**S2 Fig.** Wave propagation in 2D homogenous tissues: normal, ischemia 1a, ischemia 1b, MI, decoupled 1b and MI. (The time interval of S2 stimulation was 390ms in ischemia 1b condition and the rest was 420ms.)

**S3 Fig.** Wave propagation in the 2D tissue where ischemia 1a, decoupled ischemia 1b, and decoupled MI distributed horizontally using the S1-S2 protocol when the S2 stimulus was applied in the (A) lower left or (B) upper left corner.

**S4 Fig.** (A) (i) APD distribution in the fifth stimulation in the 2D tissue where ischemia 1a, decoupled 1b, and decoupled MI distributed circularly with a pacing cycle of 420ms. (ii) The maximum APD difference between each cell and its neighbors in the 2D tissue. (B) (i) APD of all cells along the border (line L1) in the 2D tissue. (ii) APD of all cells along the line L2 (with a radius of 350) and L3 (with a radius of 450) in the 2D tissue.

**S5 Fig.** Wave propagation in the 3D ventricular tissue (A) with ischemia 1a, (B) with ischemia 1b, or (C) with MI areas (with stimulus interval 310ms and 240ms, respectively).

**S6 Fig.** (A) (i) Wave propagation in the 2D tissue where ischemia 1a, decoupled 1b, and decoupled MI distributed horizontally and (ii) VWs when the leftmost stimulation was applied using the S1-S2 protocol before and after ischemia 1b area was replaced with ischemia 1a area. **S7 Fig.** (A) Wave propagation in the 2D tissue where ischemia 1a, 1b, and MI distributed horizontally with gradient distribution of all parameters and (B) action potentials of points P1, P2, and P3 when the leftmost stimulation was applied with a pacing cycle of 290ms using the dynamic stimulation protocol.

