## Supporting information for "Mechanisms of ventricular arrhythmias elicited by coexistence of multiple electrophysiological remodeling in ischemia: a simulation study"

Supplement

**Stimulation protocols**

Stimulation protocols in Fig 2 and S1 Fig.

The standard S1-S2 protocol was used in Fig 2 and S1 Fig with a stimulation strength of −86.2 pA/pF and a stimulus duration of 1 ms. In the S1-S2 protocol, 100 S1 stimuli with the interval of 1000 ms were applied for reaching a steady state.

Stimulation protocols in Fig 3 and S2 Fig.

The standard S1-S2 protocol was used in Fig 3 and S2 Fig with a stimulation strength of −120 pA/pF and a stimulus duration of 3 ms. In the S1-S2 protocol, five S1 stimuli with the interval of 1000 ms were applied on the leftmost three columns of nodes before S2 stimulus, which ensures the tissue reaching a steady state. S2 stimulus was applied in the lower left corner with the size of 300×300 cells in Fig 3B and S2 Fig. S2 stimulus was applied in the (Dark Red) lower left corner or (Orange) upper left corner in Fig 3C.

Stimulation protocols in Fig 4 and S3 Fig.

The standard S1-S2 protocol was used in Fig 4 and S3 Fig with a stimulation strength of −120 pA/pF and a stimulus duration of 3 ms. In the S1-S2 protocol, five S1 stimuli with the interval of 1000 ms were applied on the leftmost three columns of nodes before S2 stimulus, which ensures the tissue reaching a steady state. S2 stimulus was applied in the lower left and upper left corner with the size of 300×300 cells in Fig 4 and S3 Fig.

Stimulation protocols in Fig 5.

The standard S1-S2 protocol was used in Fig 5A with a stimulation strength of −120 pA/pF and a stimulus duration of 3 ms. In the S1-S2 protocol, five S1 stimuli with the interval of 1000 ms were applied on the leftmost three columns of nodes before S2 stimulus, which ensures the tissue reaching a steady state. S2 stimulus was applied in the leftmost three columns with the size of 3×600 cells.

The dynamic protocol was used in Fig 5B with a stimulation strength of −120 pA/pF and a stimulus duration of 3 ms. In the dynamic protocol, stimuli were applied on the leftmost three columns with the size of 3×600 cells with a pacing cycle of 250ms.

Stimulation protocols in Fig 6 and S7 Fig.

The dynamic protocol was used in Fig 6 and S7 Fig with a stimulation strength of −120 pA/pF and a stimulus duration of 3 ms. In the dynamic protocol, stimuli were applied on the leftmost three columns with the size of 3×600 cells with pacing cycles of 250ms in Fig 6 and 290ms in S7 Fig.

Stimulation protocols in Fig 7, Fig 8, and S4 Fig.

The dynamic protocol was used in Fig 7, Fig 8, and S4 Fig with a stimulation strength of −120 pA/pF and a stimulus duration of 3 ms. In the dynamic protocol, stimuli were applied on a circular sector at the upper left corner with a radius of 5 nodes with a pacing cycle of 420ms.

Stimulation protocols in Fig 9 and S5 Fig.

In the 3D ventricular tissue as shown in Fig 9 and S5 Fig, three stimuli were applied on a small cubic tissue with the size of 10×10×10 nodes in intramyocardial region with a stimulation strength of −120 pA/pF and a stimulus duration of 2 ms.

Stimulation protocols in Fig 10 and S6 Fig.

The standard S1-S2 protocol was used in Fig 10 and S6 Fig with a stimulation strength of −120 pA/pF and a stimulus duration of 3 ms. In the S1-S2 protocol, five S1 stimuli with the interval of 1000 ms were applied on the leftmost three columns of nodes before S2 stimulus, which ensures the tissue reaching a steady state. S2 stimulus was applied in the leftmost three columns with the size of 3×600 cells.

**Gradient distribution of all ion currents and ion concentration**

Decrease percentage of I_Na_ was set as gradient distribution from 11.3% (ischemia 1a) to 62% (MI). Increase percentage of I_NaL_ was set as gradient distribution from 150% (ischemia 1a) to 100% (MI). Decrease percentage of I_CaL_ was set as gradient distribution from 20% (ischemia 1a) to 50% (ischemia 1b) and from 50% (ischemia 1b) to 38% (MI). Decrease percentage of I_to_ was set as gradient distribution from 50% (ischemia 1a) to 63% (MI). [ATP]_i_ was set as gradient decrease from 6.8mM (ischemia 1a) to 4mM (MI). [K^+^]_o_ was set as gradient increase from 6mM (ischemia 1a) to 8mM (MI).

Decrease percentage of I_NaCa_ was set as gradient distribution from 100% (ischemia 1a) to 40% (ischemia 1b) and from 40% (ischemia 1b) to 100% (MI). Decrease percentage of I_NaCa_ was set as gradient distribution from 100% (ischemia 1a) to 54% (ischemia 1b) and from 54% (ischemia 1b) to 100% (MI). Decrease percentage of I_Kr_ was set as gradient distribution from 100% (ischemia 1a) to 70% (MI). Decrease percentage of I_Kr_ was set as gradient distribution from 21.9% (ischemia 1a) to 80% (MI). Increase percentage of I_bCa_ was set as gradient distribution from 100% (ischemia 1a) to 130% (ischemia 1b) and from 130% (ischemia 1b) to 100% (MI). Decrease percentage of I_rel_ was set as gradient distribution from 100% (ischemia 1a) to 35% (ischemia 1b) and from 35% (ischemia 1b) to 100% (MI). Decrease percentage of I_up_ was set as gradient distribution from 100% (ischemia 1a) to 29% (ischemia 1b) and from 29% (ischemia 1b) to 100% (MI).

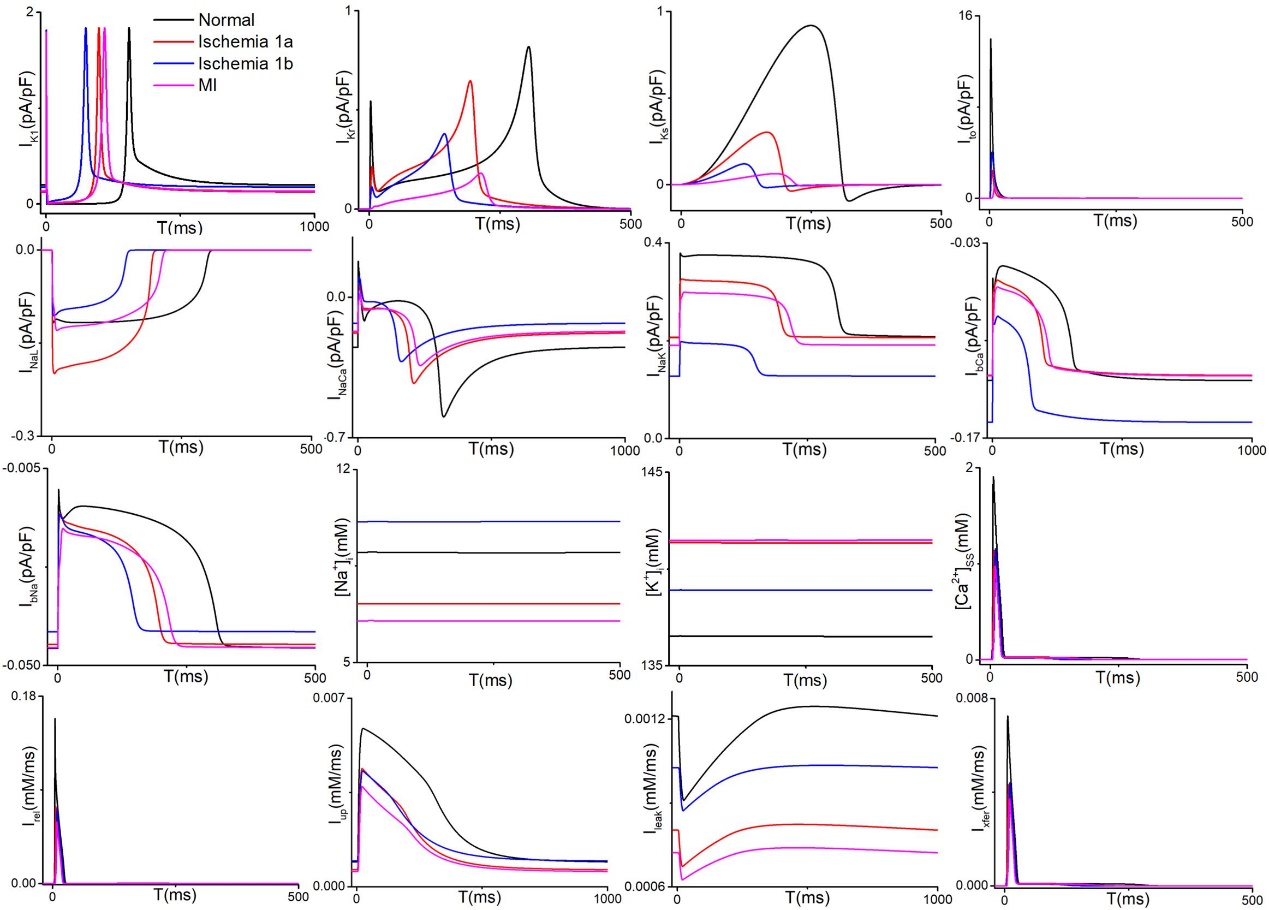

S1 Fig. The variations of electrophysiological characteristics of other ion currents and concentration of single cells in normal, ischemia1a, ischemia1b, and MI conditions.

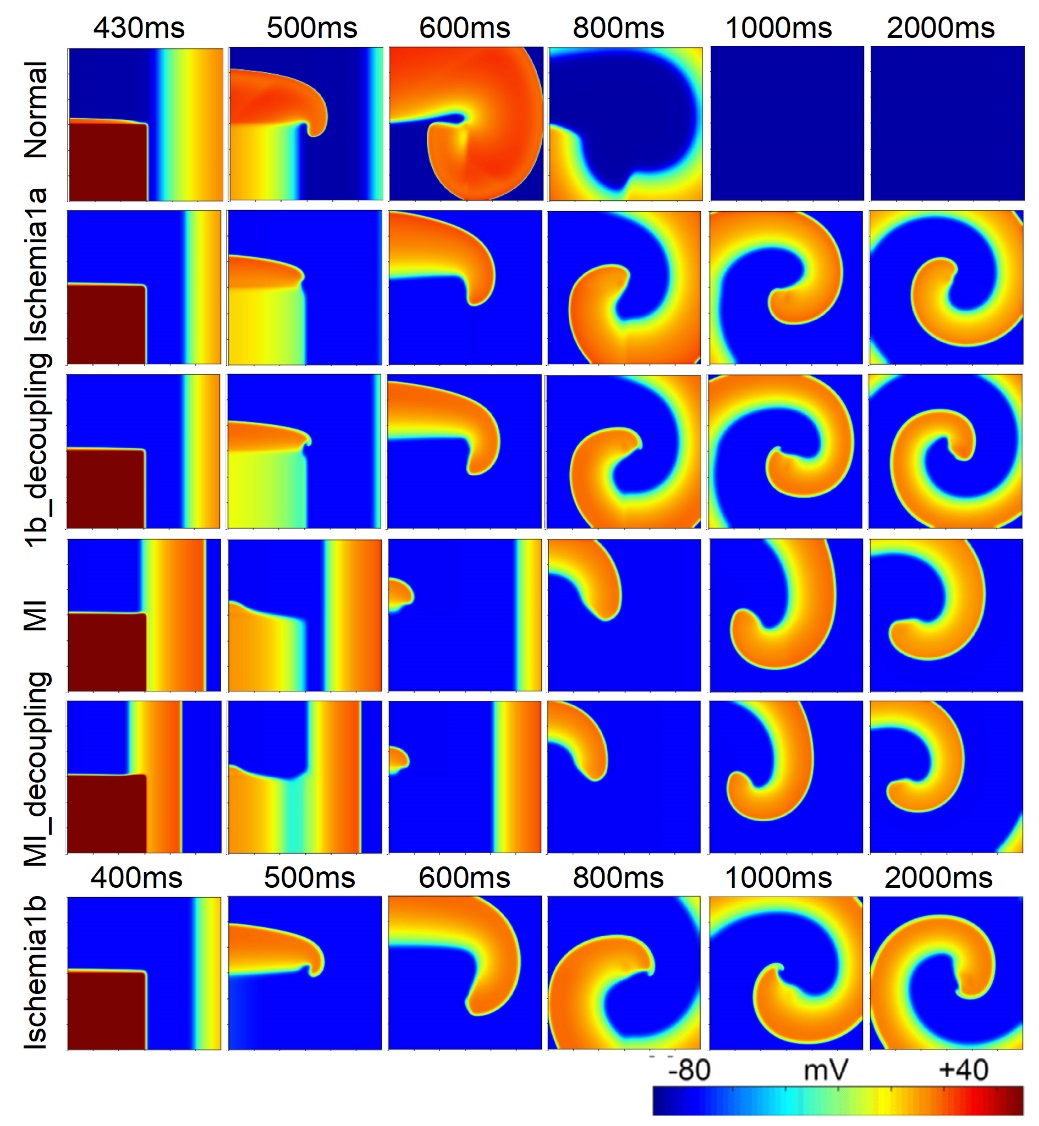

S2 Fig. Wave propagation in 2D homogenous tissues: normal, ischemia 1a, ischemia 1b, MI, decoupled 1b and MI. (The time interval of S2 stimulation was 390ms in ischemia 1b condition and the rest was 420ms.)

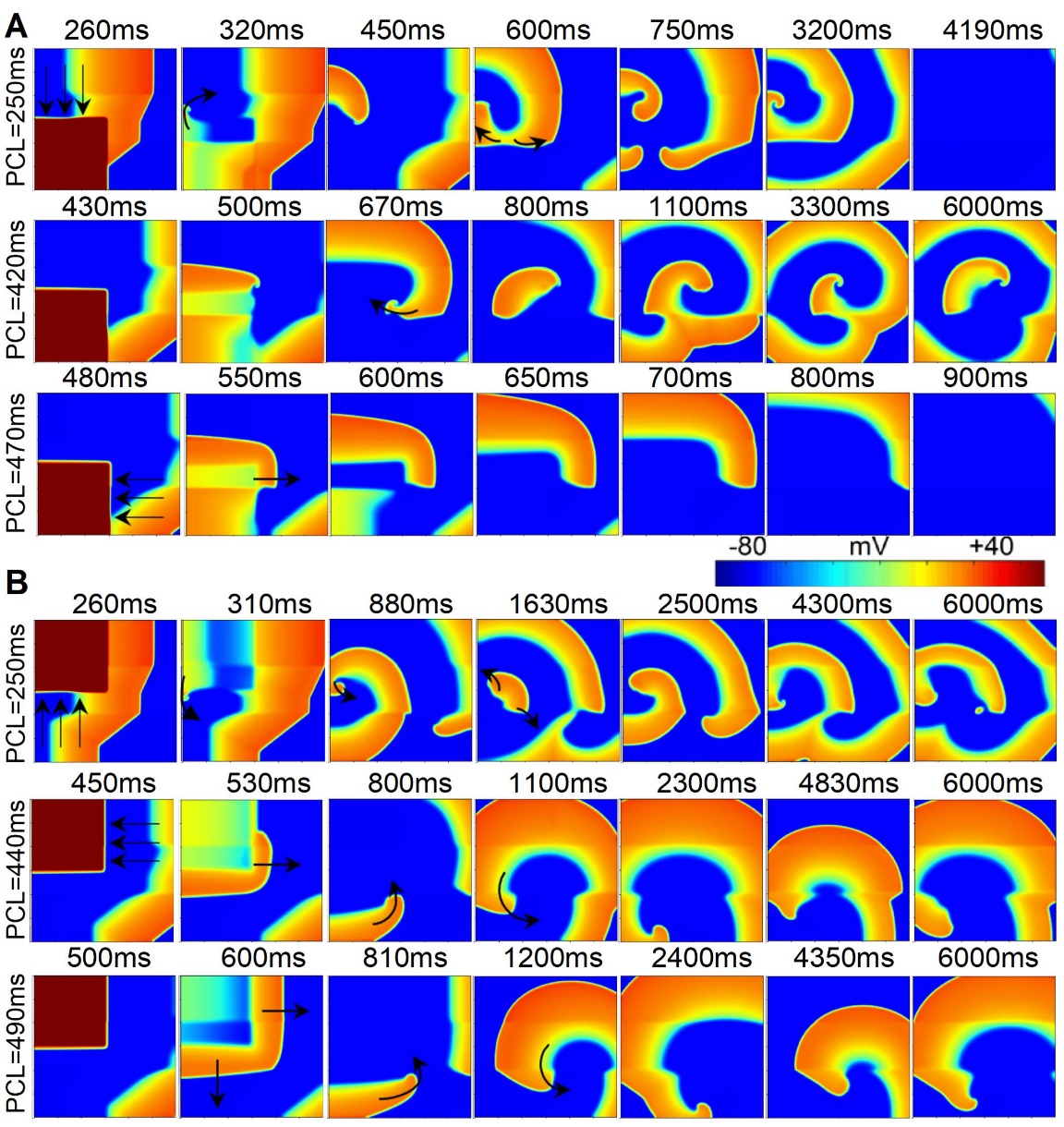

S3 Fig. Wave propagation in the 2D tissue where ischemia 1a, decoupled ischemia 1b, and decoupled MI distributed horizontally using the S1-S2 protocol when the S2 stimulus was applied in the (A) lower left or (B) upper left corner.

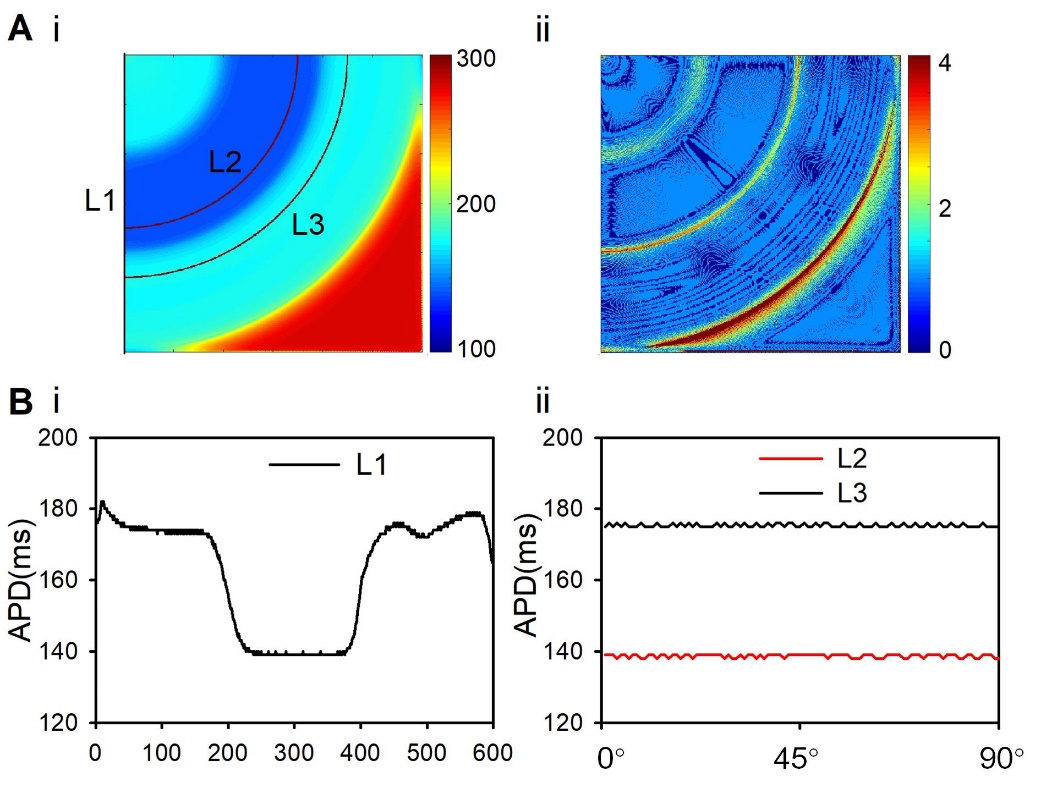

S4 Fig. (A) (i) APD distribution in the fifth stimulation in the 2D tissue where ischemia 1a, decoupled 1b, and decoupled MI distributed circularly with a pacing cycle of 420ms. (ii) The maximum APD difference between each cell and its neighbors in the 2D tissue. (B) (i) APD of all cells along the border (line L1) in the 2D tissue. (ii) APD of all cells along the line L2 (with a radius of 350) and L3 (with a radius of 450) in the 2D tissue.

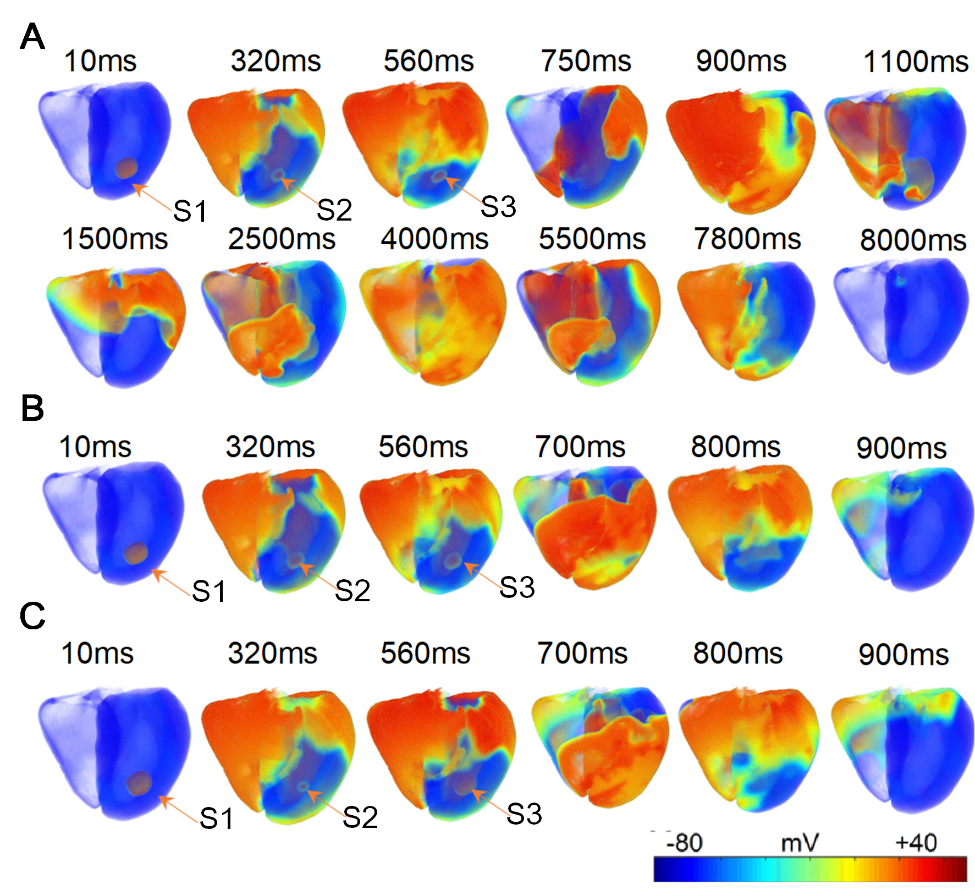

S5 Fig. Wave propagation in the 3D ventricular tissue (A) with ischemia 1a, (B) with ischemia 1b, or (C) with MI areas (with stimulus interval 310ms and 240ms, respectively).

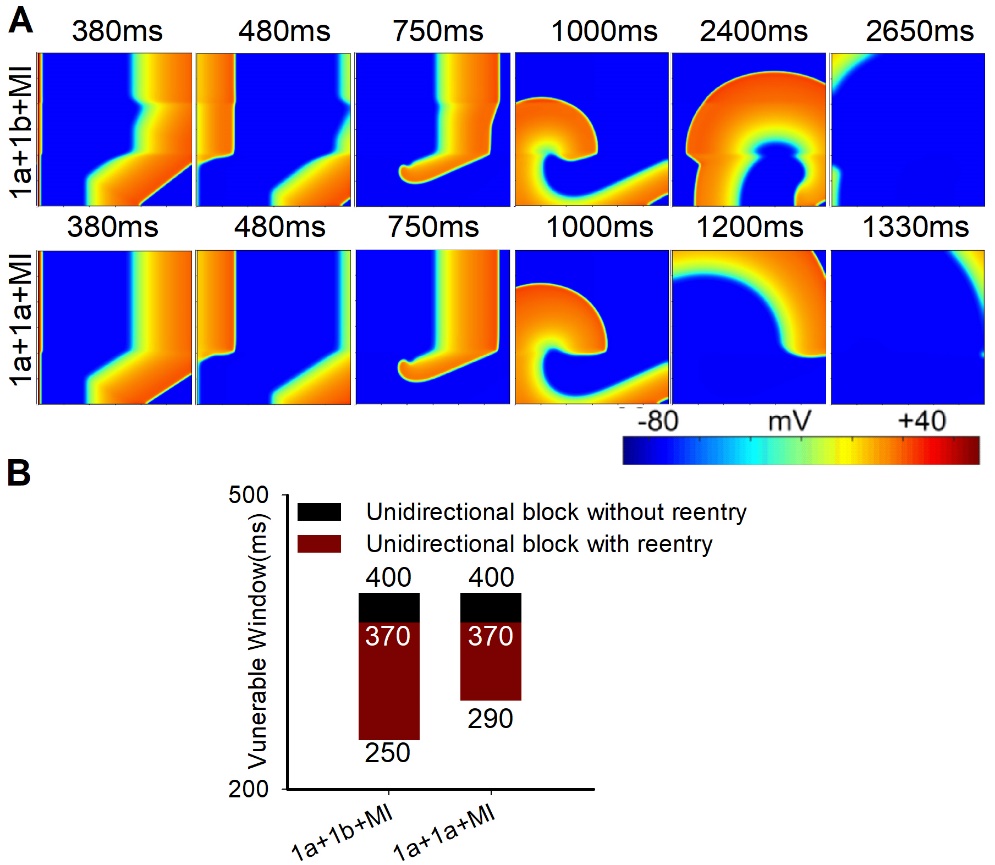

S6 Fig. (A) (i) Wave propagation in the 2D tissue where ischemia 1a, decoupled 1b, and decoupled MI distributed horizontally and (ii) VWs when the leftmost stimulation was applied using the S1-S2 protocol before and after ischemia 1b area was replaced with ischemia 1a area.

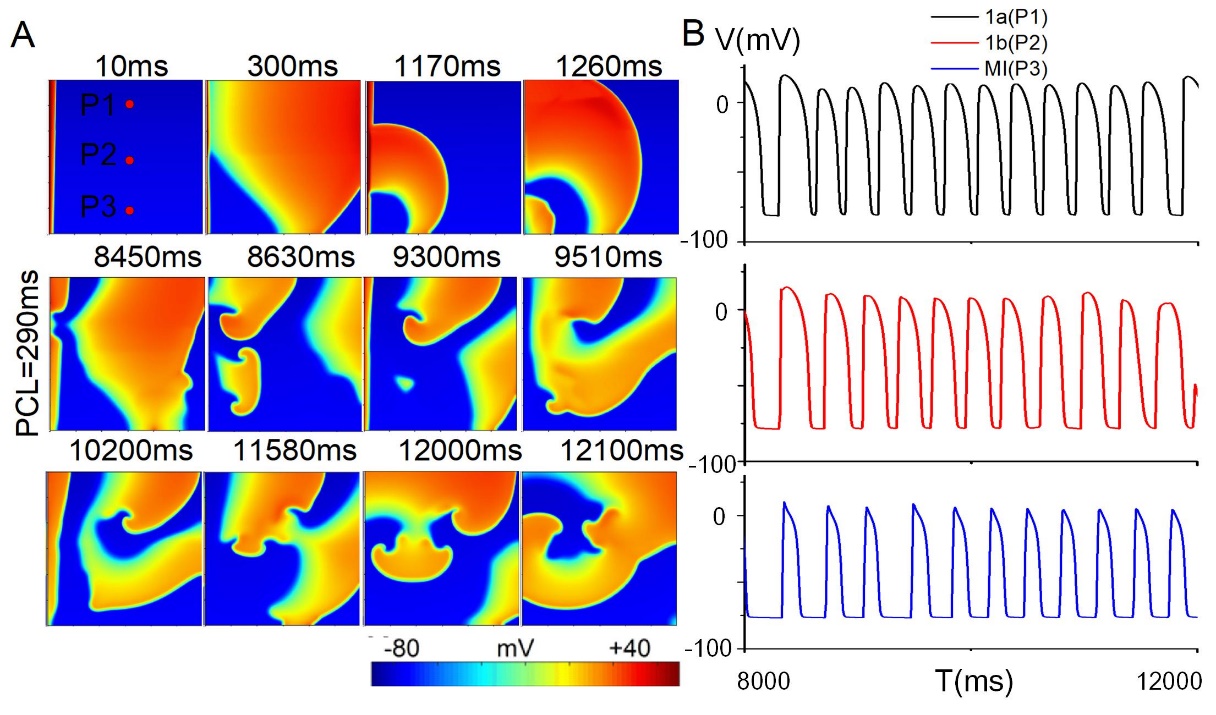

S7 Fig. (A) Wave propagation in the 2D tissue where ischemia 1a, 1b, and MI distributed horizontally with gradient distribution of all parameters and (B) action potentials of points P1, P2, and P3 when the leftmost stimulation was applied with a pacing cycle of 290ms using the dynamic stimulation protocol.
